# The SARS-CoV-2 accessory protein Orf3a is not an ion channel, but does interact with trafficking proteins

**DOI:** 10.1101/2022.09.02.506428

**Authors:** Alexandria N. Miller, Patrick R. Houlihan, Ella Matamala, Deny Cabezas-Bratesco, Gi Young Lee, Ben Cristofori-Armstrong, Tanya L. Dilan, Silvia Sanchez-Martinez, Doreen Matthies, Rui Yan, Zhiheng Yu, Dejian Ren, Sebastian E. Brauchi, David E. Clapham

**Affiliations:** Janelia Research Campus, Howard Hughes Medical Institute, Ashburn, VA, 20147, USA; Physiology Institute, Universidad Austral de Chile, and Millennium Nucleus of Ion Channel-Associated Diseases (MiNICAD), Valdivia, 511-0566, Chile; Department of Biology, University of Pennsylvania, Philadelphia, PA 19104, USA; Department of Microbiology and Immunology, Cornell University, Ithaca, NY 14853, USA; Center for Advanced Imaging, University of Queensland, St. Lucia, QLD 4072, Australia; Molecular Biology Department, University of Wyoming, Laramie, WY, 82071, USA; Unit on Structural Biology, Division of Basic and Translational Biophysics, Eunice Kennedy Shriver, National Institute of Child Health and Human Development, National Institutes of Health, Bethesda, MD, 20892, USA

## Abstract

The severe acute respiratory syndrome associated coronavirus 2 (SARS-CoV-2) and SARS-CoV-1 accessory protein Orf3a colocalizes with markers of the plasma membrane, endocytic pathway, and Golgi apparatus. Some reports have led to annotation of both Orf3a proteins as a viroporin. Here we show that neither SARS-CoV-2 nor SARS-CoV-1 form functional ion conducting pores and that the conductances measured are common contaminants in overexpression and with high levels of protein in reconstitution studies. Cryo-EM structures of both SARS-CoV-2 and SARS-CoV-1 Orf3a display a narrow constriction and the presence of a basic aqueous vestibule, which would not favor cation permeation. We observe enrichment of the late endosomal marker Rab7 upon SARS-CoV-2 Orf3a overexpression, and co-immunoprecipitation with VPS39. Interestingly, SARS-CoV-1 Orf3a does not cause the same cellular phenotype as SARS-CoV-2 Orf3a and does not interact with VPS39. To explain this difference, we find that a divergent, unstructured loop of SARS-CoV-2 Orf3a facilitates its binding with VPS39, a HOPS complex tethering protein involved in late endosome and autophagosome fusion with lysosomes. We suggest that the added loop enhances SARS-CoV-2 Orf3a ability to co-opt host cellular trafficking mechanisms for viral exit or host immune evasion.

## Introduction

The β-coronavirus, <u>S</u>evere <u>A</u>cute <u>R</u>espiratory <u>S</u>yndrome <u>Co</u>rona<u>V</u>irus <u>2</u> (SARS-CoV-2) has among the largest genomes of any RNA virus^1, 2^, encoding for 29 proteins. The majority of these proteins are ‘Non-Structural Proteins’, NSPS, mediate viral RNA replication while the ‘structural proteins’ are components of the virion.^1, 2^ The third group of functionally enigmatic ‘accessory’ proteins likely bolster SARS-CoV-2 replication or facilitate evasion from the host’s innate immune system. One of these proteins is Open reading frame 3a (Orf3a), a membrane protein that is annotated as a putative viroporin based on previous work and its similarity to SARS-CoV-1, also claimed to be a viroporin.^3–6^ In this paper we determine the structures of SARS-CoV-2 and SARS-CoV-1 Orf3a, measure their functions, and examine the differences between the CoV-2 and CoV-1 proteins.

Viroporins are viral membrane proteins that commonly form weakly- or non-selective oligomeric holes in surface plasma membranes or intracellular organellar membranes.^7, 8^ SARS-CoV-1 Orf3a was postulated to be a viroporin based on its membrane localization, oligomerization, and C-terminal sequence similarities to a calcium ATPase from *Plasmodium falciparum* and an outer-membrane porin from *Shewanella oneidensis.*^5, 9^ Recording K^+^-selective currents from SARS-CoV-1 Orf3a-expressing *Xenopus laevis* oocytes were interpreted to mean that SARS-CoV-1 Orf3a was a viroporin.^5^ This current was inhibited 5-50 µM emodin, a natural compound suggested as a potential SARS-CoV-1 antiviral^5, 10^, and by 10 mM BaCl_2_, a common K^+^ channel blocker. Whole-cell patch clamp of SARS-CoV-1 expressing HEK293 cells exhibited non-selective ion currents blocked by BaCl_2_ that were also attributed to SARS-CoV-1 Orf3a.^6^ Although these data were used to support the SARS-CoV-1 Orf3a viroporin hypothesis, further work to identify its pore-lining residues is lacking.^5, 6^ One study pursued this question by testing SARS-CoV-1 Orf3a pore mutants in artificial bilayers.^11^ However, this method is prone to background channel contamination and has been recently questioned.^12, 13^

The putative viroporin activity of Orf3a has not been convincingly linked to its reported functions during SARS-CoV-1 infection. Comparison of viral infections between wild-type and Orf3a-deficient SARS-CoV-1 strains demonstrates that Orf3a promotes host cell death and causes intracellular vesicle formation and Golgi fragmentation.^14^ In addition, during SARS-CoV-1 infection, Orf3a stimulates the production of mature IL-1β, a marker of NLRP3 inflammasome activation.^15^ Although viroporins from other viruses have been shown to trigger apoptosis or innate immune cell activation, many other viral hijacking mechanisms can produce these phenotypes.^7, 8^ The lack of compelling evidence connecting the contributions of Orf3a to SARS-CoV-1 pathogenesis with its purported viroporin activity raises the question of whether its channel function is physiologically required and if SARS-CoV-1 Orf3a is a *bona fide* viroporin.

Given its similarity to the SARS-CoV-1 homolog, we asked if SARS-CoV-2 Orf3a is a viroporin. While pursuing this, several groups reported conflicting findings that support or oppose the SARS-CoV-2 Orf3a viroporin hypothesis.^3, 4, 16^ We reasoned that the discrepancy could be due to common experimental problems that arise when characterizing putative ion channels, leading to its misidentification as a viroporin.^12, 17, 18^ Additionally, SARS-CoV-2 Orf3a viroporin activity has not been studied in mammalian cells, which represents a more native environment for functional studies.^3, 4, 16^ We performed a comprehensive structural and functional investigation of SARS-CoV-2 Orf3a. We surveyed the subcellular localization of overexpressed SARS-CoV-2 Orf3a in HEK293 cells and observed co-localization with markers of the plasma membrane and the endocytic pathway, as previously reported.^19–22^ We then made extensive efforts to record SARS-CoV-2 Orf3a attributable cation currents at the plasma membrane and in endo-lysosomes of HEK293 cells. We also attempted to measure SARS-CoV-2 Orf3a currents at the plasma membrane of *Xenopus* oocytes, and in a reconstituted system. In all cell lines and reconstituted systems tested, we did not measure currents attributable to SARS-CoV-2 Orf3a. We explored this further by resolving three high-resolution cryo-EM structures of SARS-CoV-2 Orf3a under different conditions, varying the lipid composition and the scaffold protein used for nanodisc assembly. All structures were captured in the same conformational state, displaying a constriction within the transmembrane region and the presence of a basic aqueous vestibule, which would not favor cation permeation. We were also unable to recapitulate the published SARS-CoV-1 Orf3a recordings from *Xenopus* oocytes and HEK293 cells. Finally, we show that the SARS-CoV-1 Orf3a cryo-EM structure mirrors the overall architecture and structural features seen in the SARS-CoV-2 homolog. From our data, we conclude that SARS-CoV-1 and SARS-CoV-2 Orf3a are not viroporins.

What is the function of SARS-CoV-2 Orf3a and how may it contribute to SARS-CoV-2 pathogenicity? We observe Rab7 enrichment, a marker for late endosomes upon SARS-CoV-2 Orf3a overexpression, and co-immunoprecipitation with the host protein, VPS39. In contrast, SARS-CoV-1 Orf3a does not cause the same cellular phenotype and does not interact with VPS39. We identified an unstructured, cytosolic loop unique to SARS-CoV-2 Orf3a that contributes to its interaction with VPS39. Our data is in agreement with previous work showing that SARS-CoV-2 Orf3a binds VPS39, a HOPS complex tethering protein involved in autophagosome and late endosome fusion with lysosomes.^22–25^ Disruption of HOPS complex activity may be a mechanism to promote SARS-CoV-2 viral egress, an exit strategy recently proposed for β-coronaviruses.^19, 23^

## Results

### SARS-CoV-2 Orf3a colocalizes with subcellular markers of the plasma membrane and the endocytic pathway

SARS-CoV-1 Orf3a is enriched in the Golgi apparatus and plasma membrane, but is also present in late endosomes, lysosomes, and the perinuclear region when expressed in epithelial, fibroblast and osteosarcoma immortalized cell lines.^6, 9, 14, 26–29^ To determine whether SARS-CoV-2 Orf3a displays a similar subcellular distribution as SARS-CoV-1 Orf3a, we generated doxycycline-inducible HEK293 cell lines which stably expressed SARS-CoV-2 or SARS-CoV-1 Orf3a fused to a self-labeling enzyme (SNAP or HALO tag). Cells not treated with doxycycline served as negative controls. To assess localization of SARS-CoV-2 and SARS-CoV-1 Orf3a, cells were transfected or immunostained with various subcellular markers of the endoplasmic reticulum (ER), Golgi apparatus (G), plasma membrane (PM), early endosomes (EE), late endosomes (LE), lysosomes (Lyso), and peroxisomes (PX). After 24 h expression, SARS-CoV-2 Orf3a_HALO_ colocalized with the PM marker, farnesyslated-GFP, which was further supported by total internal reflection microscopy (*Figure 1A-C, Figure 1-figure supplement 1)*. We also observe partial colocalization of SARS-CoV-2 Orf3a_HALO_ with markers of the endocytic pathway including EE (EEA, Rab5), LE (Rab7) and the Lyso (LAMP1) compartments (*Figure 1D-F, Figure 1-figure supplement 1*). Minimal SARS-CoV-2_HALO_ Orf3a is seen in all other subcellular compartments tested (*Figure 1G-I, Figure 1-figure supplement 1*) consistent with other reports.^19–22^ For SARS-CoV-1 Orf3a_HALO_, we identify a similar trend of colocalization with markers of the endocytic pathway and at the plasma membrane (*Figure 1-figure supplement 2*).

**Figure 1.**
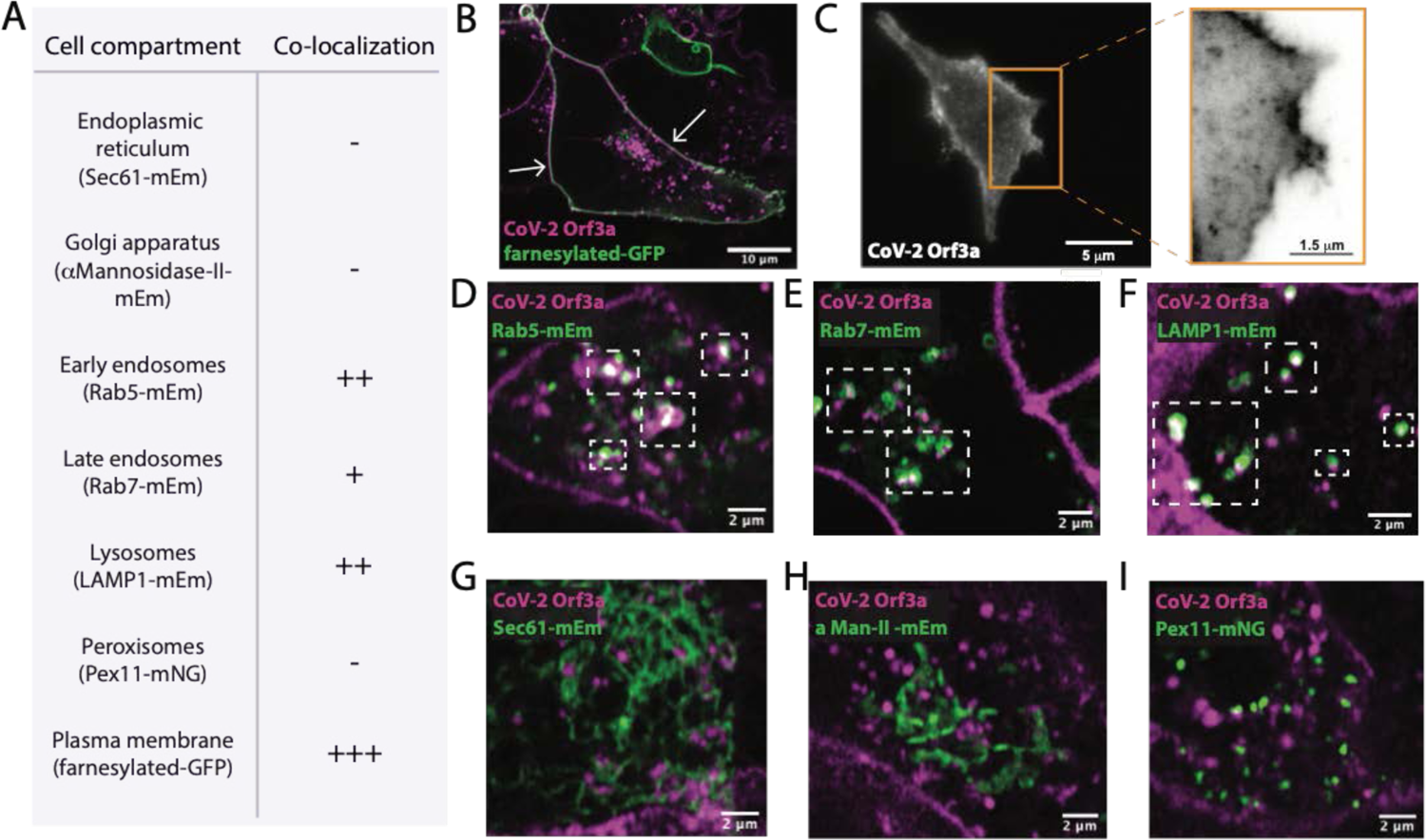
Sub-cellular localization of SARS-CoV-2 Orf3a (**A**) SARS-CoV-2 (CoV-2) Orf3a colocalizes with markers for the plasma membrane and endocytic pathway. All proteins markers used to identify cellular compartments are listed in the table in A and are transiently expressed (mEm, mEmerald; mNG, mNeonGreen) (**B**) Live-cell image of transiently expressed farnesylated-GFP (green) and CoV-2 Orf3a_HALO_ (magenta) using a HEK293 doxycycline-inducible CoV-2 Orf3a_HALO_ stable cell line. White arrows indicate co-localization. (**C**) Total Internal Reflection Fluorescence (TIRF) imaging of HEK293 cell with transient expression of CoV-2 Orf3a_SNAP_ (white). Orange box, magnification of the surface to highlight CoV-2 Orf3a_SNAP_ (black). (D-I) Live-cell image of transiently expressed (**D**) Rab5-mEmerald, (**E**) Rab7-mEmerald, (**F**) LAMP1-mEm, (**G**) Sec61-mEm, (**H**) αMannosidase-II-mEm, or (**I**) Pex11-mNG (green) with CoV-2 Orf3a_HALO_ (magenta) as described in (**B**). White boxes indicate regions of co-localization.

### SARS-CoV-2 Orf3a currents are not observed across cell or late endosome/lysosome membranes

SARS-CoV-1 Orf3a was reported to form viroporins at plasma membranes.^5, 10^ Given its sequence similarity to SARS-CoV-1 Orf3a, we asked whether SARS-CoV-2 Orf3a may also exhibit similar ion channel properties. We induced expression of SARS CoV-2 Orf3a_SNAP_ in HEK293 cells and performed whole-cell patch clamp electrophysiology. In all external cationic bath solutions that we tested, including K^+^, Na^+^, Cs^+^, N-methyl-D-glucamine (NMDG^+^), and Ca^2+^, we were not able to observe ionic current densities distinct from those measured in control cells (*Figure 2A-C*). We also performed whole-endolysosomal patch clamp electrophysiology in HEK293 cells expressing SARS-CoV-2 Orf3a_HALO_. No K^+^, Na^+^, Ca^2+^, or H^+^ currents were recorded above those measured from untransfected endolysosomal vesicles (*Figures 2D-I, Figure 2-figure supplement 1*). Our interrogation of SARS-CoV-2 Orf3a overexpressed at the PM and in the LE/Lyso of HEK293 cells suggests that that SARS CoV-2 Orf3a is not forming a cation permeable viroporin.

**Figure 2.**
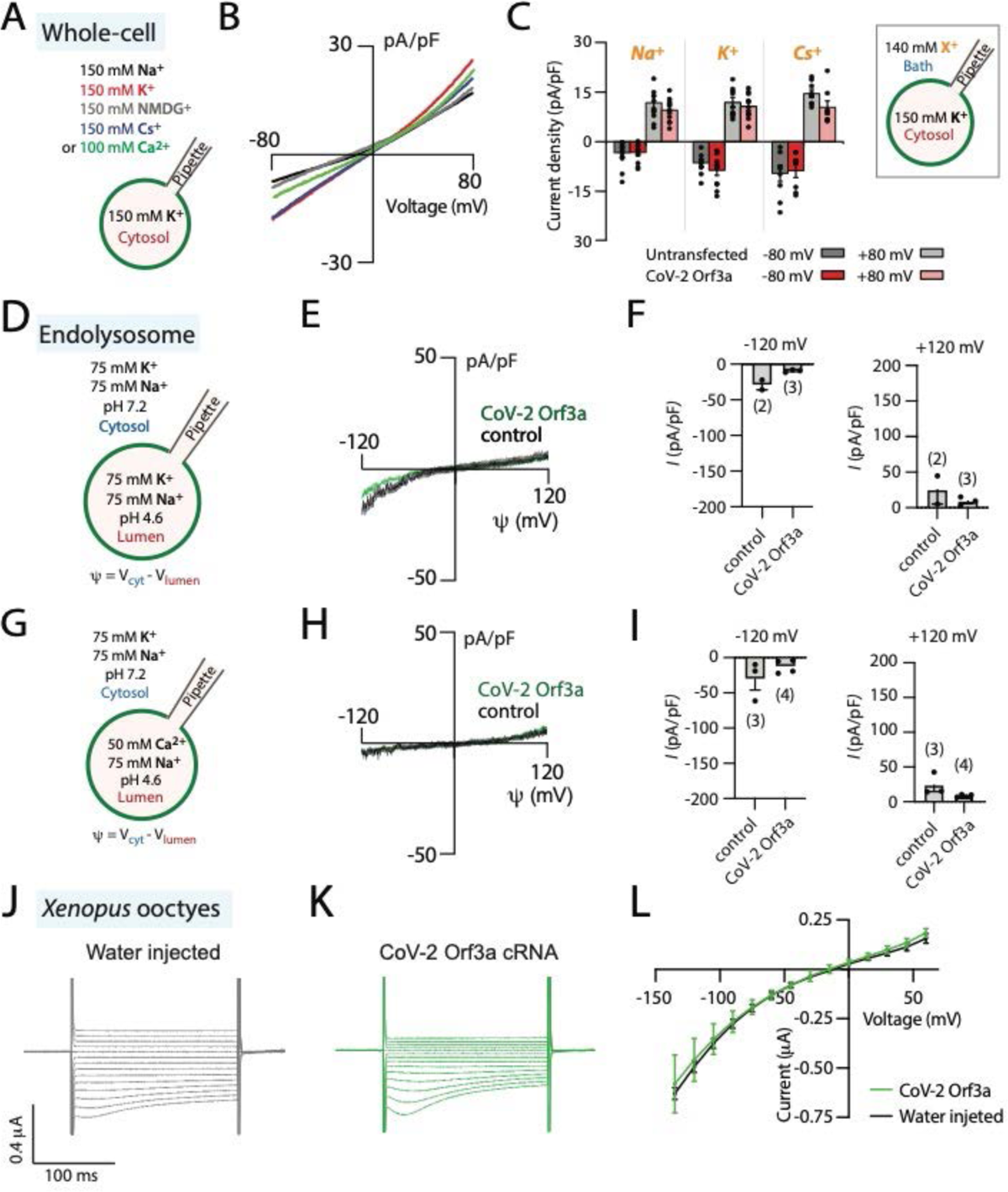
SARS-CoV-2 Orf3a is not a viroporin. (**A-C**) SARS-CoV-2 (CoV-2) Orf3a does not elicit a cation current at the plasma membrane. (**A**) Solutions used for whole-cell patch-clamp experiments. (**B**) I-V relationship for HEK293 cells expressing CoV-2 Orf3a_SNAP_ by doxycycline induction in various external cationic solutions (Na^+^, n=27; K^+^, n=5; Cs^+^, n=8; NMDG^+^, n=8; Ca^2+^, n= 5). Traces are colored based on Figure 2A (**C**) Average current density for untransfected HEK293 cells (gray bars) and cells transfected with CoV-2 Orf3a_SNAP_ (red bars) at −80 and +80 mV recorded in Na^+^ (n=11), K^+^ (n=8), and Cs^+^ (n=8) solutions. Error is represented as SEM. (**D-I**) CoV-2 Orf3a does not elicit a Na^+^, K^+^, or Ca^2+^-selective current in endolysosomes. (**D, G**) Solutions used in the endolysosomal patch clamp experiments. Cl^-^: all the bath solutions contained 150 mM Cl and pipette solutions contained 5mM Cl^-^ (**E, H**) I-V relationship for HEK293 cells expressing GFP (control, black) or CoV-2 Orf3a_HALO_ (green). (**F,I**) Average current density for control and CoV-2 Orf3a_HALO_ expressing HEK293 cells at −120 mV and +120 mV from (d,g). ns, p > 0.05 by unpaired two-tailed t tests. (J-L) CoV-2 Orf3a does not elicit a current in *Xenopus* oocytes when recorded in high K^+^ external solution. (**J-K**) Representative current traces from *Xenopus* oocytes injected with (**J**) water or (K) CoV-2 Orf3a_2x-STREP_ mRNA (20 μg). Recordings are done in high external K^+^ (96 mM KCl) that recapitulate published methods (**L**) I-V relationship for water injected (black, n=7) or CoV-2 Orf3a (green, n=7) following protocol in (**J-K**).

### SARS-CoV-2 Orf3a currents are not observed in Xenopus oocytes

We next asked whether we could mimic the conditions used to characterize SARS-CoV-1 Orf3a in *Xenopus* oocytes to investigate SARS-CoV-2 Orf3a channel activity. We generated SARS-CoV-2 Orf3a_2xSTREP_ cRNA and injected 20 ng of cRNA or water as a control into defolliculated *Xenopus* oocytes. After 48-72 hours, oocytes were recorded by two-electrode voltage clamp (TEVC) using a high potassium solution (96 mM KCl) similar to an extracellular solution that previously elicited SARS-CoV-1 Orf3a ionic currents, or standard TEVC recording solutions (ND96, 96 mM NaCl). We were not able to record SARS-CoV-2 Orf3a_2xSTREP_ ionic currents above background (*Figure 2J-L, Figure 2-figure supplement 2*), despite confirmation of PM localization of SARS-CoV-2 Orf3a_2xSTREP_ in *Xenopus* oocytes by surface biotinylation (*Figure 2-figure supplement 2*). These data further support the conclusion that SARS-CoV-2 Orf3a is not a viroporin.

### SARS-CoV-1 Orf3a currents are not observed in Xenopus oocytes or HEK293 cells

The lack of channel activity observed for SARS-CoV-2 Orf3a may reflect a functional and evolutionary distinction between SARS-CoV-2 and SARS-CoV-1 Orf3a. To explore this further, we attempted to record currents of SARS-CoV-1 Orf3a from *Xenopus* oocytes and HEK293 cells. Although we confirmed SARS-CoV-1 Orf3a PM localization in both expression systems, we recorded no currents attributable to the expressed protein (*Figure 1-figure supplement 2, Figure 2-figure supplement 2*).^5, 6, 10^ Our collective electrophysiology data indicate that neither SARS-CoV-1 nor SARS-CoV-2 Orf3a are viroporins.

### Overall three-dimensional architecture of SARS-CoV-2 Orf3a

Although our electrophysiological data suggest that SARS-CoV-2 Orf3a does not function as a viroporin at the PM or in the endocytic pathway, we sought to explore this further by determining two cryo-EM structures of SARS-CoV-2 Orf3a in nanodiscs that contain lipids which mimic each of these compartments (*Figure 3, Figure 3-figure supplement 1-3*). We first evaluated the oligomeric state of SARS-CoV-2 Orf3a_2xSTREP_ in cell membranes by chemical crosslinking and concluded that it likely assembles as a 64 kDa dimer (*Figure 3-figure supplement 2L*). Despite its challenging size for cryo-EM structural determination (< 100 kDa), we were able to resolve a nearly identical conformation of SARS-CoV-2 Orf3a in both nanodisc preparations, determined to 3.0 Å in the LE/Lyso environment or 3.4 Å in the PM environment (global RMSD 0.34, *Figure 3A, Figure 3-figure supplement 1-4*).^30^ Cryo-EM density for SARS-CoV-2 Orf3a is well-resolved between serine 40 and valine 237, whereas the electron density for the distal N- and C-termini, and a loop within the C-terminus (residues 175-180), is poor or not present. These regions are excluded from the final model (*Figure 3A, Figure 3–figure supplement 1-4*).

**Figure 3.**
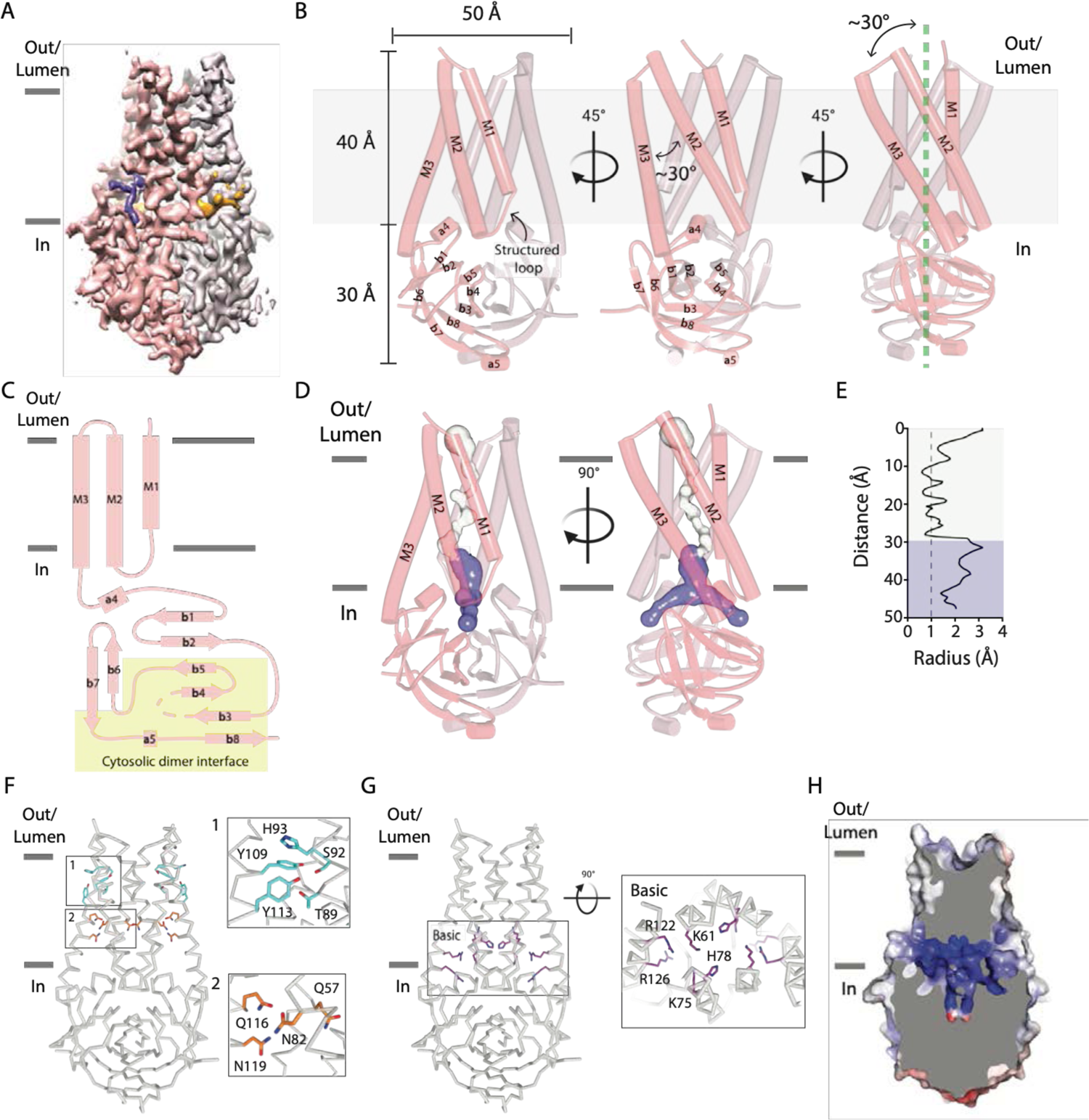
A narrow cavity detected in the SARS-CoV-2 Orf3a TM region is unlikely to conduct cations. (**A-C**). Overall architecture of SARS-CoV-2 (CoV-2) Orf3a. (**A**) Cryo-EM map of dimeric CoV-2 Orf3a (dark and light pink), with density for lipids colored (orange, purple). (**B**) Three side views of CoV-2 Orf3a depicting dimeric architecture (dark and light pink) and key structural elements. (**C**) 2D topology of CoV-2 Orf3a. The region forming the cytosolic dimer interface is shown (yellow). (**D**) Inspection of the CoV-2 Orf3a TM region for a pore, depicted as the minimal radial distance from its center to the nearest van der Waals contact (HOLE program).^73^ A region too narrow to conduct ions (white) and an aqueous vestibule (dark blue) are highlighted. (**E**) Radius of the pore (from D) as a function of the distance along the ion pathway. Dashed lines indicate the minimal radius that would permit a dehydrated ion. Blue and white colors follow (**D**). (**F**) Two layers of polar residues (1 and 2, cyan and orange) identified in the TM region, with a zoom-in of each region. (**G**) Basic residues located in the aqueous vestibule (purple) with zoom-in of the region. (**H**) Cutaway view of the CoV-2 Orf3a molecular surface is colored according to the electrostatic potential (APBS program).^74^ Coloring: blue, positive (+10 kT/e) and red, negative (−10 kT/e).

The overall architecture of SARS-CoV-2 Orf3a is a dimer and is comprised of six transmembrane (TM) helices, three provided by each subunit, with dimensions of 50 Å in diameter and 70 Å in height (*Figure 3B*). Due to the odd number of TM helices per subunit, the SARS-CoV-2 Orf3a N-terminus is positioned towards the extracellular or luminal space and its C-terminus within the cytosol, as previously reported for SARS-CoV-1 Orf3a (*Figure 3B-C*)^5, 26^. When viewed from the extracellular/luminal side, TM 1-3 are arranged in clockwise manner with pronounced tilting of TM2 and 3 (> 30°) from a line perpendicular to the membrane (*Figure 3B*). Helical tilting is likely required for TM2 and 3, which are 45 Å in length, hydrophobic, and would otherwise unfavorably protrude from the membrane. TM1 is shorter by comparison (30 Å) and extends the remainder of the lipid bilayer by a structured loop that connects with TM2. Although tilted at a similar angle in the membrane, TM2 and 3 are positioned at a 30° angle with respect to one another (*Figure 3B)*. Consequently, the combined angle between TM2-3 and the structured loop between TM1-2 create several gaps per subunit that expose the protein core to the membrane, accommodating weak to moderately resolved electron density which we attribute to lipids (*Figure 3A-B, 4A-B, Figure 4-figure supplement 1*).

**Figure 4.**
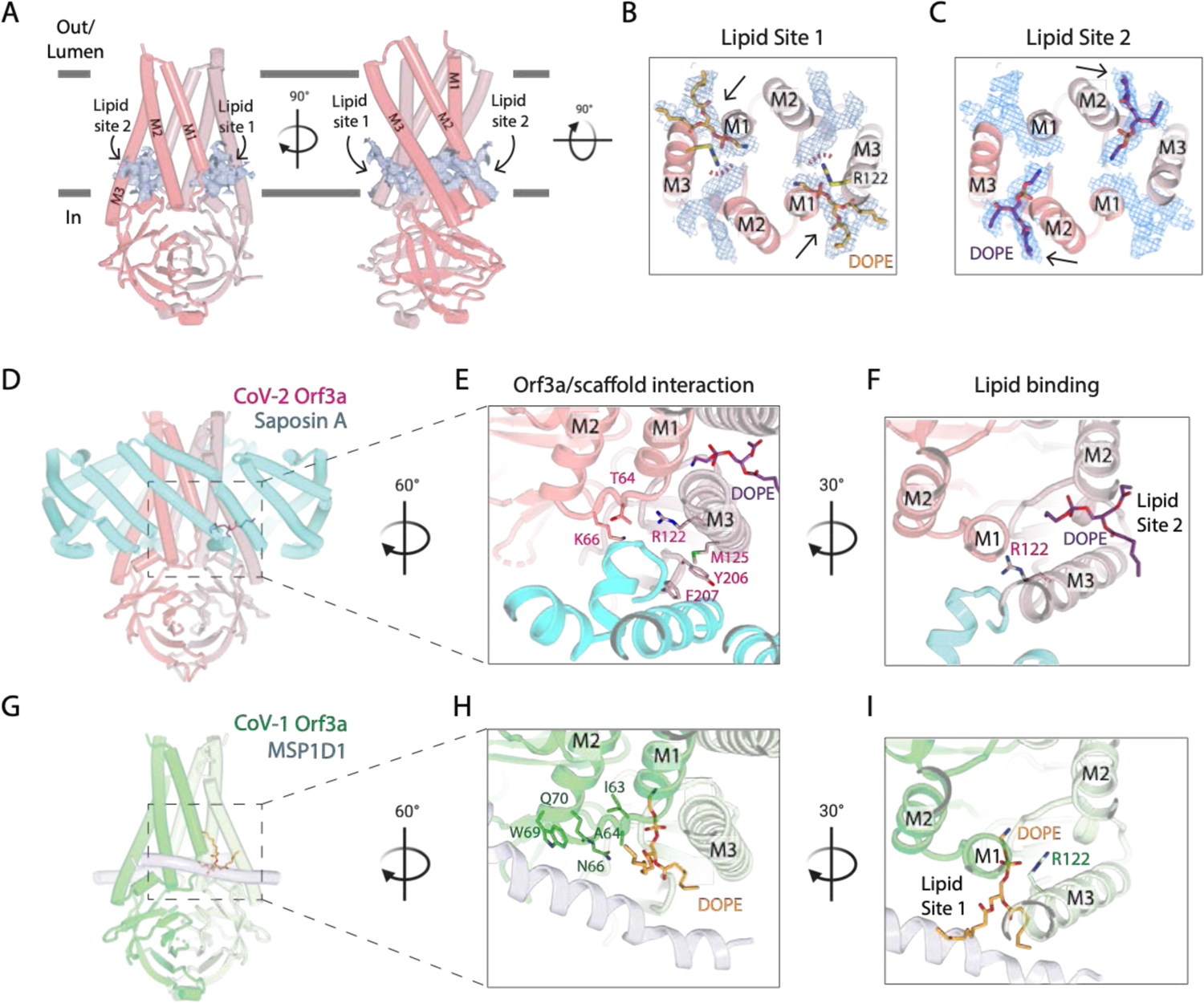
Two SARS-CoV-2 Orf3a lateral openings within the TM region are filled with density likely representing lipid sites. (**A**) Two side views of SARS-CoV-2 (CoV-2) Orf3a in LE/Lyso MSP1D1 nanodiscs highlighting two subunits (dark and light pink). Lipid densities (light blue surface and dark blue mesh, contoured at 2.5α) are identified in fenestrations between TM1 and TM3 of neighboring subunits (Lipid Site 1) and TM2 and TM3 of the same subunit (Lipid Site 2). (**B-C**) Cutaway view from the extracellular space to view lipids modeled into the density. (**B**) Two DOPE lipids (orange) are modeled into Lipid Site 1 (light blue mesh) and (c) and into Lipid Site 2 (purple). Lipid Sites 1 and 2 are likely not occupied simultaneously since the orientation of R122 (yellow) would sterically clash with DOPE in Lipid Site 2 (red dotted line). (**D-F**) CoV-2 Orf3a interacts with Saposin A. (**D**) Side view of CoV-2 Orf3a in LE/Lyso Saposin A nanodiscs highlighting two subunits (dark and light pink) and 6 molecules of Saposin A (cyan), with DOPE shown (purple). (**E**) Zoom-in from the extracellular side to highlight the CoV-2 Orf3a and Saposin A interaction. CoV-2 Orf3a residues within 5 Å from Saposin A are shown (light pink) with DOPE (purple). (**F**) Zoom-in from the extracellular side to highlight DOPE in Lipid Site 2 (purple). Note that residue R122 (light pink) is rotated 135° from the CoV-2 Orf3a LE/Lyso MSP1D1 structure (compare with Figure 4B; see also Figure 4I for direct comparison) and occludes Lipid Site 1. (**G-I**) SARS-CoV-1 (CoV-1) Orf3a interacts with MSP1D1. (**G**) Side view of CoV-1 Orf3a in LE/Lyso MSP1D1 nanodiscs highlighting two subunits (dark and light green) and two molecules of MSP1D1 (light blue), with DOPE shown (orange). (**H**) Same view as Figure 4E to highlight the CoV-1 Orf3a and MSP1D1 interaction. CoV-1 Orf3a residues within 5 Å of MSP1D1 are shown (green), with DOPE (purple) depicted. (**I**) View as in Figure 4F to highlight DOPE in Lipid Site 1 (orange). Similar to CoV-2 Orf3a reconstituted in LE/Lyso MSP1D1 nanodiscs, residue R122 (green) is positioned near and clashes with Lipid Site 2 (Figure 4b).

Extending from TM3 is a 104 amino acid structured cytosolic domain assembled from a cluster of eight β-sheets (β1-8) contributed by each SARS-CoV-2 Orf3a subunit, forming a compact and continuous molecular surface that protrudes 30 Å into the cytosol (*Figure 3B-C*). Packing of the dimer is facilitated by β3-5 and β8, as well as loops joining β2-3, β4-5 and a helical turn and loop connecting β7-8 at its base (*Figure 3B-C*). These extensive interactions along the dimer interface within the cytosolic domain (buried surface interface of 1010 Å^2^ per subunit) appear integral to the integrity of the SARS-CoV-2 Orf3a dimer (*Figure 3C*).

### A narrow cavity detected in the SARS-CoV-2 Orf3a transmembrane region likely does not represent a pore

The hallmark of all ion channels is an aqueous and often hydrophilic pore that span the TM region of the protein. To structurally evaluate whether SARS-CoV-2 Orf3a may be a viroporin that conducts K^+^ or other cations, we inspected the TM region of the protein for a channel pore that fits these criteria. Approximately two-thirds of the TM region is tightly packed in its current conformational state, with the narrowest point in this region (∼0.8 Å radius) too narrow to accommodate a dehydrated cation (*Figure 3D-E*). We then inspected the amino acid composition within this region to identify polar or charged residues which might form a hydrophilic pore if SARS-CoV-2 Orf3a was in a different conformational state. Two distinct clusters of polar residues, the first positioned towards the extracellular or luminal space (T89, S92, H93, Y109, Y113) and the second situated within the center of the membrane (Q57, N82, Q116, N119) are regions that may accommodate hydrophilic ions or molecules (*Figure 3F*).

The final approximately one-third of the TM region positioned above the cytosolic domain contains a ∼12 Å diameter aqueous vestibule which is accessible from the cytosol through two narrow portals. Although the portal could permit the movement of partially hydrated cations (1.5 Å radius), the composition of residues lining the aqueous vestibule is highly basic (K61, K75, H78, R122, R126) creating a positively charged region that would not be suitable for cations (*Figure 3D-E, G-H*). The lack of a clear and identifiable pore is inconsistent with an ion channel, even if captured in a closed or non-conductive state.

### Two lateral openings within the transmembrane region are filled with electron densities attributable to lipids

Within the SARS-CoV-2 Orf3a transmembrane region are two distinct lateral fenestrations that expose the aqueous vestibule to the lipid bilayer formed between the structured loop of TM1 and TM3 (Lipid Site 1) and between TM2 and TM3 (Lipid Site 2) (*Figure 4A-C, Figure 4-figure supplement 1*). Each fenestration is filled with tubular density attributable to lipids (*Figure 4-figure supplement 1*). We modeled a 1,2-Dioleoyl-sn-glycero-3-phosphatidylethanolamine (DOPE) molecule into Lipid Site 1 based on the size, shape, and presence of this density in both PM and LE/Lyso lipid compositions (*Figure 4B, Figure 4-figure supplement 1*). Similarly, we assigned DOPE to Lipid Site 2 (*Figure 4C, Figure 4-figure supplement 1*). A notable arginine residue (R122) located in TM3 neighbors both lipid sites and, in its current conformation, stabilizes the phospholipid phosphate group in Lipid Site 1 (*Figure 4B*). It is unlikely that both fenestrations would be occupied by DOPE at once since their headgroups are positioned too close to one another and R122 clashes with Lipid Site 2 (*Figure 4C*). DOPE bound in either lipid site would contribute to the positive electrostatic landscape of the aqueous vestibule.

### Structure of SARS-CoV-2 Orf3a in a Saposin A containing nanodisc provides additional insight into lipid fenestration binding

In both SARS-CoV-2 Orf3a cryo-EM maps, we observed low-resolution density for the membrane scaffold protein (MSP) used for nanodisc assembly (MSP1D1, *Figure 3-figure supplement 3*). MSPs typically assemble as two belts that wrap around the disk, shielding the lipid bilayer from the aqueous environment. However, we observe direct binding of the MSP1D1 with SARS-CoV-2 Orf3a, which is unusual and may inadvertently stabilize a single conformational state of the protein (*Figure 3-figure supplement 3*). To circumvent this and potentially capture a different conformational state, we substituted MSP1D1 with another scaffold protein, Saposin A (*Figure 4-figure supplement 2-3*).^31–34^ We determined the cryo-EM structure of SARS-CoV-2 Orf3a in a LE/Lyso lipid Saposin A-containing nanodisc to 2.8 Å resolution. Its overall architecture and conformational state is nearly identical to the structures of SARS-CoV-2 Orf3a in MSP1D1-containing nanodiscs, harboring a TM constriction and a basic aqueous vestibule (global RMSD 0.50, *Figure 4-figure supplement 3L*). We observe a direct interaction of Saposin A with SARS-CoV-2 Orf3a, which occurs in a similar region to where MSP1D1 is binding, but with a slightly different interface of SARS-CoV-2 Orf3a (*Figure 4D, Figure 4-figure supplement 2*). Although we were unable to identify another conformational state from this dataset, the discrete interfaces of interaction between the two scaffold proteins and SARS-CoV-2 Orf3a, with little change to the SARS-CoV-2 Orf3a structure, increases our confidence that our cryo-EM structures represent a native conformational state of the protein.

Comparison of the SARS-CoV-2 Orf3a MSP1D1 and Saposin A nanodisc structures reveals several differences which can be attributed to scaffold protein binding. The binding of the Saposin A to SARS-CoV-2 Orf3a directly occludes Lipid Site 1 and consequently, electron density is absent from this site. Instead, we observe a 135° rotation of R122 into Lipid Site 1, where its side chain is directly interacting with Saposin A, creating space for a lipid to occupy Lipid Site 2 (*Figure 4F, Figure 4-figure supplement 1*). The rotation of R122 side chain and the electron density exclusively in Lipid Site 2 supports the argument that Lipid Sites 1 and 2 are not simultaneously occupied. The presence of density in both Lipid Sites 1 and 2 in the MSP1D1-containing SARS-CoV-2 Orf3a cryo-EM maps likely represents two lipid bound states that are averaged together during the 3D reconstruction.

### The three-dimensional architecture of SARS-CoV-1 Orf3a is nearly identical to SARS-CoV-2 Orf3a

To further address the possibility of a functional and evolutionary distinction between SARS-CoV-2 and SARS-CoV-1 Orf3a, we determined the structure of SARS-CoV-1 Orf3a in a LE/Lyso-like membrane MSP1D1 nanodisc to 3.1 Å resolution and compared it to the SARS-CoV-2 homolog (*Figure 4-figure supplement 3-5)*. Its overall architecture and conformational state is nearly identical to the SARS-CoV-2 Orf3a cryo-EM structures (global RMSD 0.47, *Figure 4-figure supplement 3M*). SARS-CoV-1 Orf3a does not have an obvious pore in its current conformational state and its aqueous vestibules are also electrostatically positive (*Figure 4-figure supplement 5*). In combination with our extensive efforts to identify and reproduce SARS-CoV-1 and −2 Orf3a currents in various expression systems without success, we conclude that both Orf3a homologs are not viroporins.

Supporting the idea of two distinct lipid bound states, we observe pronounced electron density for the MSP1D1 scaffold protein in this dataset and built a model (*Figure 4-figure supplement 3*). MSP1D1 binds to SARS-CoV-1 Orf3a in a similar area as Saposin A (*Figure 4E, Figure 4-figure supplement 3*). Accordingly, density was observed in Lipid Site 1 with weak density observed in Lipid Site 2, suggesting a preferred Lipid Site 1 bound state in maps of Orf3a that contain resolved MSP1D1 density (*Figure E, G, Figure 4-figure supplement 1*). This difference between the SARS-CoV-1 and SARS-CoV-2 Orf3a MSP1D1-containing nanodisc structures likely reflects a variation in particle heterogeneity between the datasets and not a distinction between Orf3a proteins. A recently published cryo-EM map of SARS-CoV-2 Orf3a in a PM lipid MSP1D1-containing nanodisc resolved a high-resolution density for MSP1D1 and a concomitant Lipid Site 1-only bound state.^3^ It is unclear what the significance of the discrete lipid bound states may be, but points to a function of Orf3a that might directly involve the lipid bilayer.

### Macroscopic K^+^ and Cl^-^ flux is not observed in vesicle-reconstituted SARS-CoV-1 and SARS-CoV-2 Orf3a

To further evaluate whether SARS-CoV-2 Orf3a might form a viroporin, we performed both flux assays and proteoliposome patch clamp experiments with purified, vesicle-reconstituted SARS-CoV-2 Orf3a (*Figure 5*)^3, 35^. We reconstituted purified SARS-CoV-2 Orf3a_2xSTREP_ at a high (1:10, 1:25) or standard (1:100, wt:wt) protein to lipid ratio to ensure detection of transport and ionic currents, and to attempt to recapitulate published results.^3^ We did not observe any K^+^ or Cl^-^ transport using an 9-Amino-6-Chloro-2-Methoxyacridine (ACMA) fluorescence-based flux assay (*Figure 5B-C, Figure-5-figure supplement 1A-C*). However, the counter ions used in these assays could also permeate through a non-selective viroporin, eliminating the ion gradient needed to generate flux. To circumvent this, we designed a 90° light-scattering K^+^ flux assay (*Figure 5-figure supplement 1D*).^36, 37^ Compared to the vesicle control, no reduction in light scattering is observed with SARS-CoV-2 Orf3a_2xSTREP_-containing vesicles (*Figure 5-figure supplement 1E-G*).

**Figure 5.**
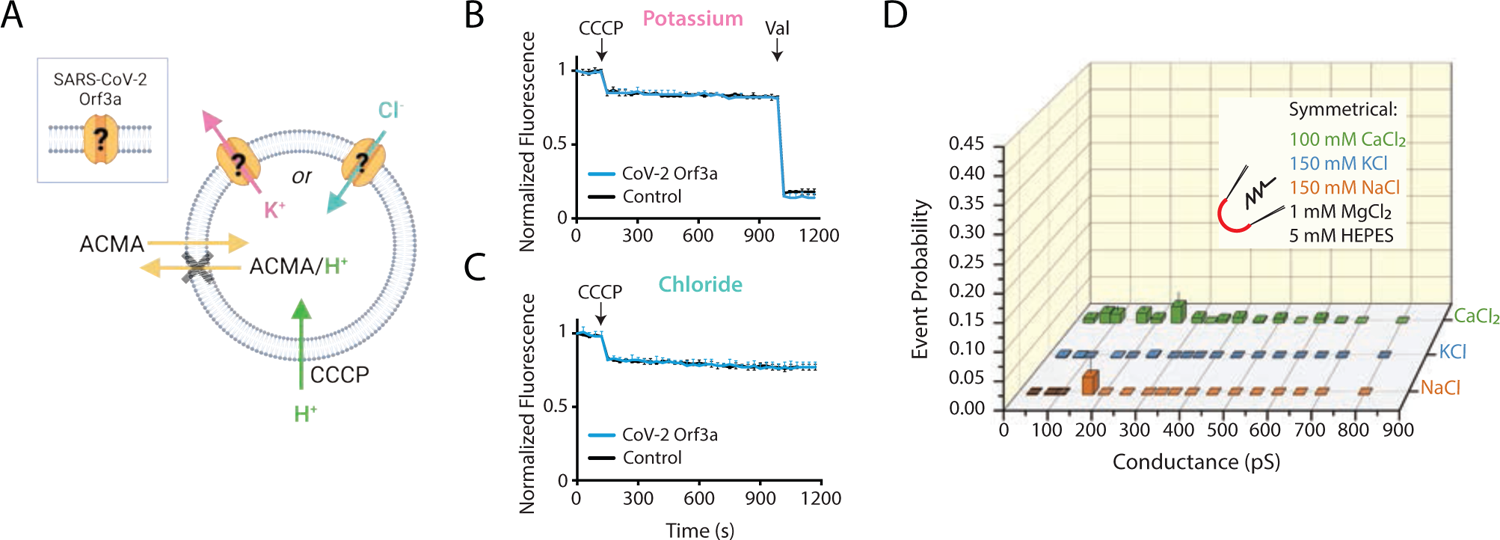
SARS-CoV-2 Orf3a does not elicit ion flux or currents in a vesicle-reconstituted system. (**A**) Schematic of the ACMA-based fluorescence flux assay.^75–78^ A K^+^ (pink) or Cl^-^ (blue) gradient is generated by reconstitution and dilution into an appropriate salt (K^+^ efflux: 150 KCl in, 150 NMDG-Cl out; Cl^-^ flux: 110 Na_2_SO_4_ in, 125 NaCl out; in mM). The addition of the protonophore, carbonyl cyanide m-chlorophenyl hydrazone (CCCP) would promote H^+^ (green) influx if CoV-2 Orf3a is a channel. ACMA is quenched and sequestered in vesicles at low pH, resulting in loss of ACMA fluorescence. Valinomycin (Val), an ionophore, is added to the end of the K^+^ flux assay to empty all vesicles. Created with Biorender.com. (**B-C**) K^+^ (n=4) (B) or Cl^-^ (n=4) (**C**) flux is not observed in CoV-2 Orf3a-reconstituted vesicles (blue) as compared with the empty vesicle control (black, n=4) using vesicles reconstituted at a 1:100 protein to lipid ratio. Error is represented as SEM. (D) Probability of observing an open event in a SARS-CoV-2 Orf3a reconstituted proteoliposome patch with vesicle reconstituted at a 1:100 protein to lipid ratio. NaCl n = 27, KCl n = 32, and CaCl_2_ n = 102. Error is represented as SEM.

### Large single-channel currents measured from vesicle-reconstituted SARS-CoV-2 Orf3a are due to transient membrane leakiness and/or channel contamination

We next asked if we could record macroscopic channel currents by proteoliposome patch clamp using vesicle-reconstituted SARS-CoV-2 Orf3a. Our attempts to record macroscopic currents were unsuccessful, yet we observed large single channel-like conductances in symmetrical K^+^, Na^+^ and Ca^2+^ from SARS-CoV-2 Orf3a_2xSTREP_ vesicles reconstituted at a high, but not standard, protein to lipid ratios (*Figure 5D, Figure 5-figure supplement 2A-C*). Although these conductance are consistent to what has been recently published, we believe that these data likely result from transient membrane leakiness/ ion channel protein contamination, and not due to SARS-CoV-2 Orf3a activity (*Figure 5-figure supplement 2B-C*).^3, 12, 18^ We examined the possibility that the single channel current measured could be contributed by an ion channel that was inadvertently co-purified with SARS-CoV-2 Orf3a. We performed mass spectrometry using proteoliposome-reconstituted SARS-CoV-2 Orf3a. The only ion channel protein that appeared on this list were the voltage-dependent anion channels 1-3 (VDAC 1-3), outer membrane mitochondrial proteins that are large-conducting, non-selective channels which can be blocked by 4,4’-Diisothiocyano-2,2’-stilbenedisulfonic Acid (DIDS) (*Figure 5-figure supplement 2D-F*).^38^ We asked if some of the currents that we record could be blocked by DIDS. After 1 min of application of the vehicle (DMSO), we observed channel block by 100 μM DIDS, which could be recovered by a washout of the inhibitor, suggesting that some of the current that we measure may be contributed by a VDAC (*Figure 5-figure supplement 2E-F*). This was a reminder that patch clamp recording detects single proteins (as single channel openings) and thus far exceeds the limits of specific protein biochemical purification.^12^ This phenomenon has been described for samples reconstituted at a high protein to lipid ratio, and is a common cause of novel protein misidentification as ion channels.^18, 35^

### SARS-CoV-2 Orf3a interacts with VPS39, a member of the HOPS complex, altering fusion of late endosomes with lysosomes

If SARS-CoV-2 Orf3a is not a viroporin, then what is its function? We turned to published proteomics datasets that have identified putative host proteins which interact with SARS-CoV-2 Orf3a to glean insight.^20, 39^ One of these host proteins, VPS39, is a member of the HOPS complex, a membrane tethering complex that facilitates LE or AP fusion with lysosomes.^25^ Defects in HOPS complex-mediated trafficking leads to the accumulation of these organelles.^40–45^ In doxycycline-treated HEK293 cells expressing SARS-CoV-2 Orf3a_HALO_, we observe an accumulation of Rab7 puncta, which is not observed in uninduced, control cells, suggesting a defect in HOPS complex-mediated vesicle fusion (*Figure 6A*).^22–24^ We then asked if a similar phenomenon is observed in the presence of SARS-CoV-1 Orf3a. In contrast, we do not observe the Rab7 puncta after doxycycline-induced expression of SARS-CoV-1 Orf3a_HALO_, suggesting an evolutionary distinction between these two homologs (*Figure 6B*).^22–24^

**Figure 6.**
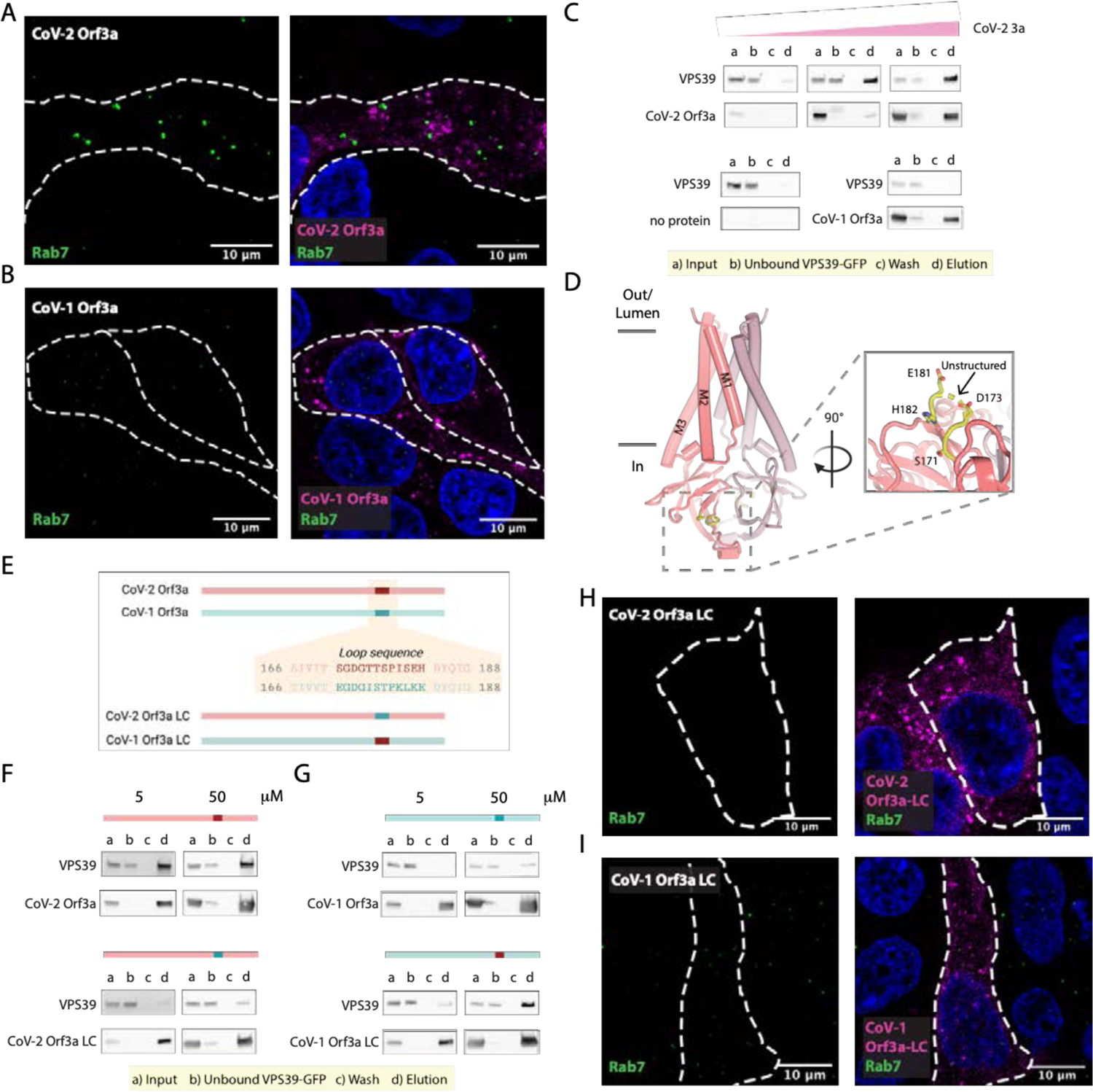
SARS-CoV-2 Orf3a, but not SARS-CoV-1 Orf3a, interacts with HOPS complex protein, VPS39. (**A-B**) Rab7 puncta (green) are abundant in HEK293 cells expressing (**A**) SARS-CoV-2 (CoV-2) Orf3a_HALO_ (magenta), but not (B) SARS-CoV-1 (CoV-1) Orf3a_HALO_ (magenta; nuclei, blue). (**C**) VPS39 interacts with CoV-2 Orf3a_2x-STREP_ in a concentration-dependent manner (top left, 0.5 μg; middle 5 μg, right, 50 μg), but does not interact with CoV-1 Orf3a_2x-STREP_ (bottom right; 50 μg). Control is Co-IP without Orf3a_2x-STREP_ added (bottom left). (**D-I**) An unstructured loop of CoV-2 Orf3a partially mediates its interaction with VPS39. (**D**) Side view of CoV-2 Orf3a structure with the subunits (dark and light pink) and unstructured loop highlighted (yellow, dotted gray box). Zoom-in of the loop from the cytosol (solid gray box) with resolved loop residues shown. (**E**) Comparison of CoV-2 Orf3a (red) and CoV-1 Orf3a (blue green) loop sequences. Orf3a wild-type (WT) and loop chimeras (LC) are color matched or swapped. Created with Biorender.com (**F**) Co-IP of CoV-2 Orf3a with VPS39 is reduced with LC and, conversely, (**G**) CoV-1 Orf3a LC interaction is enhanced with VPS39. (**H, I**) Rab7 puncta (green) are absent in HEK293 cells expressing CoV-2 Orf3a LC_HALO_ (H, magenta) or CoV-1 Orf3a LC_HALO_ (I, magenta; nuclei blue), consistent with previous reports.^23^

We next asked whether SARS-CoV-1 and SARS-CoV-2 Orf3a binds to the HOPS complex protein, VPS39. We anticipated that VPS39 likely interacts with SARS-CoV-2 Orf3a, but not SARS-CoV-1 Orf3a, based on our Rab7 accumulation data. For these co-immunoprecipitation (co-IP) experiments, we titrated increasing concentrations of purified Orf3a_2xSTREP_ with cell lysate containing overexpressed VPS39_GFP_ (*Figure 6C*). We observed an enrichment of VPS39_GFP_ that was SARS-CoV-2 Orf3a_2xSTREP_ concentration dependent (0.5-50 μg protein; *Figure 6C*), but very little to no binding was observed when performing the co-IP with all concentrations of SARS-CoV-1_2xSTREP_ Orf3a (50 μg shown; *Figure 6C*).

### An unstructured loop unique to SARS-CoV-2 Orf3a interacts with VSP39

Since we observe an interaction of VPS39 with SARS-CoV-2 Orf3a, but not SARS-CoV-1 Orf3a, we asked if we could identify the site of the protein-protein interaction. By inspecting the cytosolic domain of both Orf3a cryo-EM structures and by pinpointing the divergent regions of amino acid sequence conservation, we identified an unstructured loop between β3-4 that could be facilitating the VPS39 interaction (*Figure 6D*). We generated chimeras of SARS-CoV-2 Orf3a and SARS-CoV-1 Orf3a by swapping the 12 amino acid loop sequence between these two proteins and observe a significant reduction of interaction between SARS-CoV-2 Orf3a_2xSTREP_ loop chimera (LC) and VPS39_GFP_ (*Figure 6E, F, Figure 6-figure supplement 1*). Conversely, by substituting the SARS-CoV-2 Orf3a 12 amino acid loop into the SARS-CoV-1 Orf3a (SARS-CoV-1 Orf3a_2xSTREP_ LC), we noticed enhanced interaction with VPS39_GFP_ compared with SARS-CoV-1 Orf3a_2xSTREP_ (*Figure 6E, G, Figure 6-figure supplement 1*). However, this substitution did not fully restore the binding of VPS39 with SARS-CoV-2 Orf3a (compare with *Figure 6C and F*).

We next asked if overexpression of SARS-CoV-1 and −2 Orf3a LC could alter HOPS complex-mediated fusion, as assessed by the accumulation of Rab7-containing vesicles. Overexpression of SARS-CoV-2 Orf3a_HALO_ LC by doxycycline in HEK293 cells did not promote Rab7 accumulation, consistent with its weaker interaction with VPS39 (*Figure 6F, H*). SARS-CoV-1 Orf3a_HALO_ LC also did not promote Rab7 accumulation, as one would expect if the Orf3a and VPS39 interaction was fully rescued (*Figure 6I*). This is consistent with the co-IP experiments and suggests that the binding interface involves more than the unstructured loop (*Figure 6G, I, Figure 7A*). Overall, we observe an interaction of VPS39 that is specific to SARS-CoV-2 Orf3a, resulting in a Rab7 accumulation phenotype typical of a HOPS complex fusion defect, and that this interaction is partially mediated by a divergent, unstructured loop of SARS-CoV-2 Orf3a.

**Figure 7.**
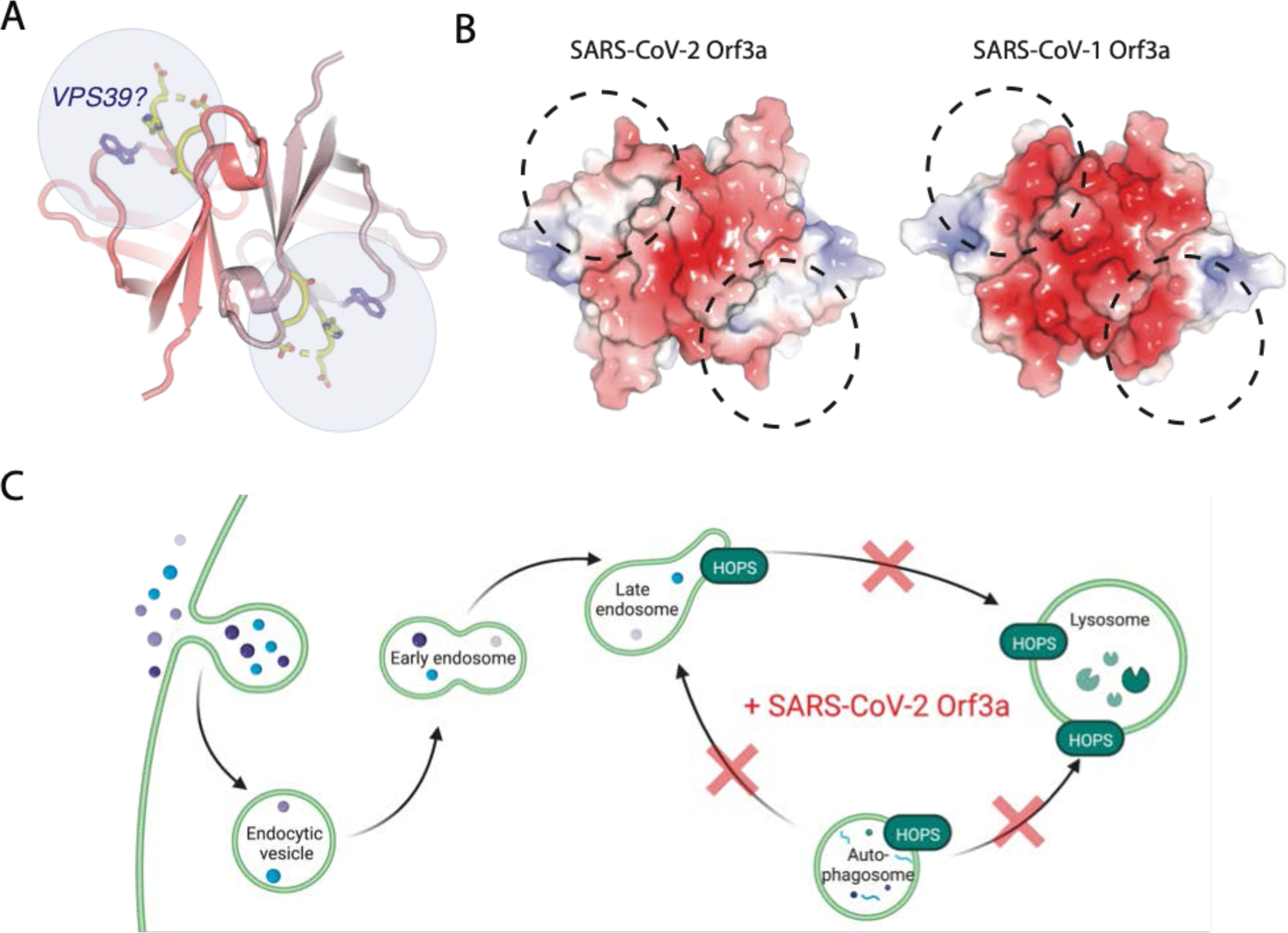
A region of VPS39 and SARS-CoV-2 Orf3a interaction. (**A**) Cytosolic view of SARS-CoV-2 Orf3a structure (dark and light pink) with the unstructured loop highlighted in yellow. W281 has also been described to mediate an interaction between SARS-CoV-2 Orf3a and VPS39 (purple).^23^ Putative VPS39 interfaces are indicated (light blue spheres) with potential stoichiometries of 1:1 or 1:2 molecules of VPS39 to SARS-CoV-2 Orf3a. (**B**) Cytosolic surface of SARS-CoV-2 and SARS-CoV-1 Orf3a colored by their electrostatic potential (APBS program): blue, positive (+5 kT/e), red, negative (−5 kT/e).^74^ The putative VPS39 interface (dotted black lines) is the same as indicated in A. (**C**) Working model of SARS-CoV-2 Orf3a dysregulation of LE and AP fusion with lysosomes. HOPS-dependent regions of the endocytic and autophagy pathways that are disrupted by SARS-CoV-2 Orf3a. Adapted from “Mutation of HOPS Complex Subunits”, by BioRender.com (2022). Retrieved from https://app.biorender.com/biorender-templates.

## Discussion

Viroporins are small viral membrane proteins that often form weakly- or non-selective pores.^7, 8^ They are widely distributed among different viral families and accordingly, their contributions to viral survival range widely from assisting with viral propagation to participating in host immune evasion.^7, 8^ Four proteins encoded by SARS-CoV-1 have been proposed to function as viroporins - E protein, Orf3a, Orf8a and Orf10.^5, 17, 46, 47^ To demonstrate that a novel protein unequivocally forms a viroporin requires the identification of its pore-lining residues. This is ideally done by introducing a pore-lining cysteine that can be modified by a methanethiosulfonate regent, which when added, partially blocks the pore or alters ion channel selectivity.^48, 49^ This has yet to be demonstrated for SARS-CoV-1 or SARS-CoV-2 Orf3a.

We asked if Orf3a from SARS-CoV-2 is a *bona fide* viroporin and took a comprehensive approach to investigate its function. We first assessed sub-cellular localization of SARS-CoV-2 Orf3a to determine where it may be functioning in mammalian cells and observed its enrichment at the PM and in the endocytic pathway. We then performed whole-cell patch clamp in HEK293 cells and endo-lysosomal patch clamp from HEK293 cells, in numerous cationic conditions. We also tested SARS-CoV-2 Orf3a at the PM of *Xenopus* oocytes and in reconstituted systems using purified SARS-CoV-2 Orf3a. We were unable to record currents that we can attribute to SARS-CoV-2 Orf3a.

While performing these studies, several other groups published electrophysiology data to either support or oppose the SARS-CoV-2 Orf3a viroporin hypothesis.^4, 16^ Our results from *Xenopus* oocytes are consistent with Grant and Lester.^16^ We propose that the reported SARS-CoV-2 Orf3a current observed by Toft-Bertelsen *et al.* could be contributed by the calcium-activated chloride channel TMEM16A, a known background channel in *Xenopus* oocytes, also suggested by others.^4, 17, 50^ Similar to Kern *et al.*, we were able to measure multiple large single channel-like conductances from vesicle-reconstituted SARS-CoV-2 Orf3a in K^+^, Na^+^, and Ca^2+^ conditions using an atypically high protein to lipid ratio (1:10, wt:wt).^3, 35^ These channel-like conductances were not present in samples reconstituted using a standard protein to lipid mixture of 1:100 (wt:wt).^35^ The lack of macroscopic currents recorded from vesicles containing high protein amounts was concerning since we estimate that ∼1000 putative channels/μm^2^ should be present.^51, 52^ We suspect that these channel-like conductances reflect spontaneous leak due to the high protein amounts in the sample, or possibly a contaminating channel such as VDAC1.^18^ We did not observe K^+^ or Cl^-^ flux from any of our reconstituted sample tested in several flux assays, further indicating that the channel-like conductances are likely spurious.

To attempt to capture SARS-CoV-2 Orf3a in different conformational states, we determined three cryo-EM structures of SARS-CoV-2 Orf3a in lipid environments mimicking the PM and LE/Lyso and in a nanodisc containing a different membrane scaffold protein. We observe one nearly identical state in all SARS-CoV-2 Orf3a sample preparations, consistent with the published cryo-EM structure of SARS-CoV-2 Orf3a, determined at 2.1 Å resolution.^3^ We do not observe a membrane-spanning pore wide enough to permit the movement of dehydrated cations. Furthermore, the positive electrostatic potential within the aqueous vestibule would not favor cation permeation.

We also attempted to repeat published data by investigating SARS-CoV-1 Orf3a viroporin activity in both HEK293 cells and *Xenopus* oocytes.^5, 6^ We were unable to record cationic currents attributable to SARS-CoV-1 Orf3a in either system, yet we observed its localization at the PM. Similarly, we were unable to observe K^+^ flux from vesicle-reconstituted SARS-CoV-1 Orf3a. We also determined a cryo-EM structure of SARS-CoV-1 Orf3a in an LE/Lyso MSP1D1-containing nanodisc and found its overall architecture indistinguishable from the SARS-CoV-2 homolog. From our comprehensive evaluation of both SARS-CoV-1 and SARS-CoV-2 Orf3a in multiple cell lines and reconstituted systems, and the conservation between the cryo-EM structures, we conclude that SARS-CoV-1 and −2 Orf3a are not viroporins.

Although a wide, membrane-spanning pore is not evident from the SARS-CoV-1 and SARS-CoV-2 Orf3a structures, we do observe density in our cryo-EM maps that we attribute to lipids. By comparing our four cryo-EM maps – three SARS-CoV-2 Orf3a samples reconstituted into nanodiscs containing PM- or LE/lyso-like lipid compositions and either MSP1D1 or Saposin A protein scaffolds, and one SARS-CoV-1 Orf3a sample reconstituted into a MSP1D1-containing nanodisc in a LE/Lyso-like environment – we detect two independent lipid binding sites likely occupied by DOPE. Although the lipids in our nanodiscs preparations are symmetrically distributed, the orientation of the two Orf3a lipid binding sites toward the cytosol and the known cellular distribution of PE on the inner leaflet of the PM and likely LE/Lyso suggests physiological relevance.^53^ We reason that Orf3a is not engaged with both molecules simultaneously due to steric clashes with R122 and between the two DOPE headgroups, as well as differences in lipid density presence or absence between the cryo-EM datasets. This is consistent with the published cryo-EM structure of SARS-CoV-2 Orf3a, in which they observe density in Lipid Site 1 and high-resolution density for the MSP1D1 scaffold protein, as we observe in our SARS-CoV-1 Orf3a map.^3^

The possibility of two discrete sites for lipid binding points to a function that involves direct interaction with lipids in the bilayer. Intracellular vesicle formation is a cellular hallmark of a SARS-CoV-1 infection, possibly contributing to the region of viral RNA replication that is proposed to provide protection from host immune responses.^14^ These intracellular vesicles are absent in Vero cells infected with SARS-CoV-1 ΔOrf3a, suggesting that Orf3a may be necessary for the formation of the vesicles.^14^ Since the membrane vesicle structures are also observed during a SARS-CoV-2 infection, it is plausible that Orf3a may be required for this process. This would suggest a function shared by SARS-CoV-1 and −2 Orf3a whereby direct interaction with the lipid bilayer promotes the formation of these vesicle structures by an unknown mechanism.

In addition to its interaction with lipids, we asked if there are host proteins that may bind to SARS-CoV-1 or −2 Orf3a to better understand its possible contributions to viral pathogenesis. We turned to published proteomics datasets and focused on VPS39, a member of the HOPS membrane tethering complex that is essential for LE or AP fusion with Lyso.^20, 25, 39^ A cellular defect common to HOPS complex mutants is an accumulation of LE and AP. We observe an enrichment of Rab7 puncta in cells overexpressing SARS-CoV-2 Orf3a_HALO_, but not SARS-CoV-1 Orf3a_HALO_. Consistent with this, VSP39 selectively interacts with SARS-CoV-2 Orf3a, and not SARS-CoV-1 Orf3a. By generating Orf3a chimeras (LC) we showed that an unstructured loop of SARS-CoV-2 Orf3a mediates binding to VPS39. Although SARS-CoV-1 Orf3a LC displays enhanced interaction with VPS39, it is not fully restored, and overexpression of the chimera is not sufficient to generate the Rab7 enrichment phenotype. Overall, our data suggests that the VPS39 interaction with Orf3a was acquired in SARS-CoV-2 and, by sequence and structural comparison between SARS-CoV-1 and SARS-CoV-2 Orf3a, that we have likely identified SARS-CoV-2 Orf3a’s novel region of interaction.

Concurrent with our cellular and biochemical studies connecting SARS-CoV-2 Orf3a overexpression with HOPS complex fusion defects, several other groups published comprehensive work that support our findings.^22–24^ In particular, Chen *et al.* surveyed the cytosolic domain of SARS-CoV-2 Orf3a to find a region of VPS39 interaction and identified a similar binding surface.^23^ Two SARS-CoV-2 Orf3a residues, S171 and W193, were identified to be important for mediating this interaction.^23^ Our Orf3a loop chimeras includes the S171 substitution and their results support our conclusion that SARS-CoV-1 Orf3a LC can partially rescue the VPS39 interaction, but it is not sufficient to restore the SARS-CoV-2 Orf3a Rab7 phenotype.^23^ Chen *et al.* demonstrate that a W193 substitution into SARS-CoV-1 Orf3a can partly restore their SARS-CoV-2 Orf3a cellular phenotype, further delineating the SARS-CoV-2 Orf3a binding surface.^23^ A comparison of the surface electrostatic potential in this putative region of VPS39 binding highlights an acidic region of SARS-CoV-1 Orf3a which is neutral on the SARS-CoV-2 Orf3a surface (*Figure 7*). We propose that the unstructured loop, W193 substitution, and the neutral charge in this region contributes to promoting the SARS-CoV-2 Orf3a and VPS39 interaction.

How does the acquired interaction between SARS-CoV-2 Orf3a and VPS39 contribute to viral pathogenesis? Several recent studies have implicated the endocytic pathway as the primary mode of viral egress for several β-coronaviruses, including SARS-CoV-2, and that Orf3a may mediate this process.^19, 23^ One possibility is that the SARS-CoV-2 Orf3a and VPS39 interaction promotes viral exit by perturbing forward endocytic trafficking through disrupting the HOPS complex-mediated fusion of LE with Lyso (*Figure 7C*). This could be due to 1) Orf3a sequestering VPS39 from the HOPS complex or, 2) Orf3a interacting with the HOPS complex through VPS39, an important distinction that has yet to be elucidated. A second, related possibility is that the SARS-CoV-2 Orf3a and VPS39 interaction promotes viral egress by interacting with known membrane tethering, fusion and trafficking complexes that facilitate Lyso movement to the cell periphery. In particular, VPS39 knockdown in SARS-CoV-2 Orf3a overexpressing cells reduces the localization of LAMP1 vesicles near the PM, and concomitantly diminishes recruitment of protein complexes involved in Lyso PM trafficking.^23^ These data suggest that VPS39 is necessary for SARS-CoV-2 Orf3a-mediated Lyso PM trafficking and implicates the recruitment of the HOPS complex by Orf3a in this process.^23^

A third possibility is the SARS-CoV-2 Orf3a and VPS39 interaction prevents HOPS-mediated AP-Lyso fusion, which we and others have observed (Miller and Clapham, not shown, *Figure 7C*).^22, 24^ Autophagy is a intracellular surveillance process that targets damaged cellular or foreign materials such as viruses for lysosomal degradation. Many viruses hijack host cell autophagy to prevent its degradation and promote survival. Overall, the acquired SARS-CoV-2 Orf3a and VPS39 interaction may function to assist with SARS-CoV-2 exit and host intracellular immune evasion. The molecular details of the host cell response to SARS-CoV-2 Orf3a need to be examined during viral infection to fully elucidate the contributions of Orf3a to SARS-CoV-2 cell physiology and pathogenesis.

## Materials and Methods

### Antibodies and dyes

SnapTag and HaloTag Janelia Fluor dyes were obtained from the Janelia Research Campus.^54^ Rabbit monoclonal antibodies against EEA1 and against LAMP1 were from Cell Signaling Technologies (Catalog #: 3288 and 9091; RRID: AB_2096811 and AB_2687579). Mouse monoclonal antibody against Rab7 (Rab7-117) was from Sigma Aldrich (Catalog #: R8779; RRID: AB_609910). Rabbit polyclonal antibody against TGN46 was from Novus Biologicals (Catalog #: NBP1-49643; RRID: AB_10011762). Rabbit monoclonal antibody against GFP was from Abcam (Catalog #: ab32146, RRID: AB_732717). Mouse monoclonal antibody against the Strep tag (StrepMAB-Classic) was from IBA Lifesciences (Catalog #: 2-1507-001; RRID: AB_513133). Hoechst 33342 dye was from Thermo Fisher (Catalog #: H3570). Alexa Fluor 488 goat anti-mouse and goat anti-rabbit antibodies were from Thermo Fisher (Catalog # A32723 and A32731; RRID: AB_2633275 and AB_2633280). HRP-conjugated mouse and rabbit antibodies were from Jackson ImmunoResearch Laboratories Inc. (Catalog # 115-035-174 and 111-035-152; RRID AB_2338512 and AB_2337936).

### Molecular biology

Codon-optimized SARS-CoV-1 Orf3a was synthesized (GenScript) and codon-optimized. SARS-CoV-2 Orf3a was a generous gift from David Gordon and Nevan Krogan.^39^ Human VPS39 was received from the MGC collection (Horizon Discovery). Full-length SARS CoV-1 Orf3a (Uniprot P59632) and SARS-CoV-2 Orf3a (Uniprot P0DTC3) were cloned into pcDNA3.1 for transient mammalian expression, a modified Piggyback vector for Tet-inducible expression, XLone, and a modified pEZT-BM vector for expression using the BacMam system.^55, 56^ Both constructs were sub-cloned using NheI and NotI restriction sites into pcDNA3.1 and pEZT-BM vectors, or MhuI and SpeI restriction sites into the XLone vector. Constructs contained a C-terminal SNAP, HALO, or GFP moiety for cell imaging, or a Twin-Strep affinity purification tag that could be removed by HRV-3C protease. Full-length human VSP39 (Uniprot Q96JC1-2) was sub-cloned using NheI and NotI restriction sites into a modified pEZT-BM vector.^55, 56^ This construct contained a C-terminal GFP tag that could be removed by TEV protease. For cRNA generation, pcDNA3.1-containing SARS-CoV-1 Orf3a_STREP_ and SARS-CoV-2 Orf3a_STREP_ were linearized with the *XmaI* restriction enzyme. Linearized DNA was used as template for *in vitro* RNA synthesis (mMESSAGE mMACHINE™ T7 Transcription Kit; Thermo Fisher).

### Generation of HEK293T stable cell lines

HEK293T cell lines (ATCC) were cultured in DMEM (ATCC) supplemented with 10% tetracycline negative fetal bovine serum (FBS, Gemini Bio Products) at 37 °C with 5% CO_2_. For generation of stable lines, cells were detached using 0.25% Trypsin-EDTA (Thermo Fisher) and seeded at a density of 2×10^5^ and allowed to recover overnight. The following day, cells were transfected at a density of ∼2.5 ×10^5^ using either Lipofectamine 2000 (HEK293T cells) with 2 ug of each Orf3a XLone constructs and 0.8 ug of the hyperactive piggyBac transposase. After 24 h, cells that had successfully integrated Orf3a were selected with 10-30 ug/mL of blasticidin-HCl (Gemini Bio Products). Cells were passaged x2 before expanding and flash-cooling. Doxycycline-inducible expression was tested in each cell line using 3 different concentrations (0.1-1 μg/mL) of doxycycline hyclate (Sigma-Aldrich) and a no-doxycycline control to evaluate Orf3a expression leak. The cell line with the highest expression and minimal leak were selected for experiments (SARS-CoV-2 Orf3a_SNAP_, SARS-CoV-2 Orf3a_HALO_ LC and SARS-CoV-1 Orf3a_HALO_ LC) or were sorted by flow cytometry (see below) to identify a population of cells that expressed Orf3a at a high concentration based on fluorescence intensity (SARS-CoV-2 Orf3a_HALO_ and SARS-CoV1 Orf3a_HALO_).

### Flow cytometry

Cells were trypsinized, pelleted, and then resuspended in 500 µl PBS and sorted into tubes or 96-well plates with a Sony SH800 cell sorter. The gating strategy to determine the SARS-CoV-2 Orf3a_HALO_ and SARS-CoV1 Orf3a_HALO_ expressing cell populations consisted of control cell samples not incubated with Janelia Fluor®635 dye, control cells incubated with Janelia Fluor®635 dye and doxycycline-treated cells incubated with Janelia Fluor®635 dye. Samples were gated based on forward and backscatter and sorted for Janelia Fluor®635 dye events in the top 3–4% of Cy5-A channel fluorescence (Cy5: 638 nm laser excitation, 665 nm with a 30 nm bandpass emission).

### Co-localization and immuno-staining experiments

35 mm (Cellvis) or 8-well (Thermo Fisher) glass bottom dishes were pre-coated with 0.1 mg/mL poly-D lysine (Thermo Fisher) for 1 h at 37°C. Cells were seeded at a density of 1-1.5×10^5^ cells and allowed to recover overnight. For subcellular marker overexpression experiments, cells were transfected at a density of ∼2 ×10^5^ cells using Lipofectamine 2000 (Thermo Fisher) with 0.5-1 μg/μL of plasmid DNA containing Sec61-mEmerald, αMannosidase II-mEmerald, Rab5-mEmerald, Rab7-mEmerald, Lamp1-mEmerald or Pex11-mNeonGreen. Orf3a expression was induced with 100 ng/mL doxycycline after 24 h, and JF635 HALO ligand was added the following day for 1-2 h at 37°C. Cells were washed twice before being switched to Live Cell Imaging Solution (Thermo Fisher).

For immunostaining of sub-cellular markers, Orf3a expression was induced with 100ng/mL doxycycline. The following day, cells were incubated with JF635 HALO ligand 1-2 h 37°C. Cells were then fixed with 4% paraformaldehyde (Electron Microscopy Sciences) for 30 min at room temperature, rinsed x3 with PBS + 0.01% sodium azide (PBS-A). Cells were permeabilized with PBS + 0.1 % Triton-X100 for 30 min on a shaker, followed by a rinse with PBS and block with BlockAid blocking solution (Thermo Fisher) for 0.5 −1 h. Primary antibody was added using the following dilutions in BlockAid buffer: 1:200 EEA1 rabbit monoclonal, 1:500 Rab7 mouse monoclonal, 1:100 LAMP1 rabbit monoclonal, and 1:200 TGN46 rabbit monoclonal. Samples were incubated for 1 h at room temperature. Cells were washed x3 with PBS-A. Secondary antibodies were added using the following dilutions in BlockAid buffer: 1:1000 Alexa Fluor 488 goat-anti mouse or 1:1000 Alexa Fluor 488 goat anti-rabbit, and 1:5000 Hoescht 33342 stain. Samples incubated for 1 h at room temperature, washed and stored in PBS-A.

Images were acquired using a Zeiss 880 Laser Scanning Confocal microscope equipped with a Plan-Apochromat 63x oil objective, 40x multi-immersion LD LCI Plan-Apochromat objective and a 20x air Plan-Apochromat objective. Data acquisition used ZEN Black software (Zeiss Instruments). Hoechst 33342 dye was excited by 405 nm laser light and the spectral detector set to 409-481 nm. Alexa 488 dye was excited by 488 nm laser light and the spectral detector set to 490-545 nm. JF635 HALO ligand was excited with 633 nm light and the spectral detector set to 642-755 nm. The spectral detector was only used for non-Airyscan confocal scanning imaging sessions. Data analysis was performed in Image J. For cell images of Rab7 puncta, a median filter was applied to the confocal slices to remove background Rab7 staining. Data processing of Airyscan confocal scanning images used auto settings for a 3D Z-Stack in ZEN Black (Zeiss Instruments).

### Total internal reflection microscopy

Cells were imaged using an inverted Nikon Eclipse TiS microscope main body and through-the-objective TIRF mode. A 488 nm solid-state laser (Coherent, Santa Clara, CA) was used for excitation. Excitation was controlled by a mechanical shutter (Uniblitz, VA Inc., Rochester, NY). The laser beam was focused to the backplane of a high-numerical aperture objective (60X, N.A. 1.49, oil) by a combination of focusing lenses. Fluorescence emission was collected by an Orca Flash 4.0 sCMOS camera (Hamamatsu Photonics, Japan), after passing through an emission filter for the acquisition wavelength band of interest (540/40 nm, Semrock, US).

### Whole-cell electrophysiology

Doxycycline-inducible HEK293T cells, expressing Orf3a_SNAP_ from SARS-CoV-1 or SARS-CoV-2, were cultured in DMEM supplied with 10% FBS (Gibco). 48 h before electrophysiological recordings, cells where incubated in media with 100 ng/µL doxycycline, 500 nM JF505 SNAP ligand or vehicle. 1-3 h before recording, cells where rinsed x3 with PBS and then maintained in fresh recording media. Whole-cell currents were obtained from transiently transfected HEK293T cells. Gigaseals were formed using 1- to 3-MΩ borosilicate pipettes (Warner Instruments). Whole-cell voltage clamp was performed and the macroscopic currents acquired at 10 kHz and filtered at 5 kHz using an A-M Systems 2400 patch clamp amplifier. Series resistance and cell capacitance were compensated via amplifier controls. Current-voltage relationships were studied using voltage ramps or steps protocols, with sweeps every 2 s. Currents are presented in pA/pF. Recordings were digitized using a Digidata 1320A (Molecular Devices, LLC, California). The analysis was performed using Clampfit 10.3 (Molecular Devices, LLC, California).

For voltage ramps protocols performed in *Figure 2A-B*, internal solution contained 150 mM KCl, 1 mM MgCl_2_, 5 mM HEPES, pH 7.4, and 5mM EGTA. External solution used for monovalent cations were: 150 mM XCl (X = Na^+^, K^+^, Cs^+^ or NMDG^+^), 5 mM HEPES, pH 7.4, 1mM MgCl_2_, and 1mM CaCl_2_, or the divalent Ca^2+^: 100 mM CaCl_2_, 5 mM HEPES and 1 mM MgCl_2_ at pH 7.4. For voltage steps protocols presented in *Figure 2C* and *Figure 2-figure supplement 3G-I*, internal solution were 147 mM KCl, 3 mM KOH, 1 mM MgCl_2_ and 10 mM HEPES. External solution used were: 140 mM XCl (X = Na^+^, K^+^ or Cs^+^), 10 mM HEPES, pH 7.4, 2 mM MgCl_2_, 2 mM CaCl_2_ and 8 mM glucose.

### Endolysosomal patch-clamp

HEK293T cells were cultured in DMEM (Thermo Fisher Scientific) supplemented with 10% FBS (R&D Systems) and 1× penicillin–streptomycin (Thermo Fisher Scientific) at 37°C with 5% CO_2_. HEK293T cells were plated on 12-mm poly-l-lysine-coated coverslips in 24 well plate, and ORF3a-HALO was transfected using PolyJet (SignaGen). The medium was replaced with fresh DMEM containing 10% FBS and 1× penicillin–streptomycin after 6 h of transfection, followed by 1 µM vacuolin to enlarge endolysosomes. Whole-endolysosomal recordings were performed as previously described with an Axopatch 200-B amplifier and Digidata 1440A controlled by pClamp and Clampfit (Molecular Devices).^57, 58^ The recordings were done in 24–32 h after transfection. For recordings presented in *Figure 2D-I*, the bath solution contained (in mM) 75 KCl, 75 NaCl, 1 MgCl_2_, 0.25 CaCl_2_, 1 EGTA, 10 HEPES (pH 7.2). The pipette solution contained (in mM) 75 NaOH, 70 KOH, 5 KCl, 150 MSA, 1 MgCl_2_, 1 CaCl_2_, 10 MES (pH 4.6) (*Figure 2D-F*) or 50 CaOH_2_, 70 NaOH, 5 NaCl, 150 MSA, 1 MgCl_2_, 1 CaCl_2_, 10 MES (pH 4.6) (*Figure 2G-I*). For recordings presented in *Figure 2-figure supplement A-C*, the bath and pipette solution contained (in mM) 150 NMDG, 1 HCl, 10 HEPES, 10 MES (pH 4 or pH 7 by MSA).

### Two-electrode voltage-clamp

Defolliculated *Xenopus laevis* oocytes were purchased from Ecocyte Bioscience, injected with either water or ∼20 ng RNA per oocyte, and stored at 17 °C in ND96 buffer (96 mM NaCl, 2 mM KCl, 5 mM HEPES, 1 mM MgCl_2_ and 1.8 mM CaCl_2_, pH 7.5 with NaOH) supplemented with 50 μg/ml gentamycin and 1% fetal bovine serum. Recordings were performed 2–3 days after injection in ND96 or high K^+^ buffer (2 mM NaCl, 96 mM KCl, 5 mM HEPES, 1 mM MgCl_2_ and 1.8 mM CaCl_2_, pH 7.5 with KOH) at room temperature using a Warner OC-725C Oocyte Clamp amplifier (Warner Instrument Corp, USA) with a ∼500 μL recording chamber. Recording microelectrodes were pulled from borosilicate glass using a P-97 Puller System (Sutter Instrument) with resistances of 0.2–0.5 MΩ when filled with 3 M KCl. Data were acquired using the pCLAMP 10 software (Molecular Devices) and a Digidata 1440A digitizer (Molecular Devices), filtered at 0.5 kHz and digitized at 10 kHz. Oocytes were subject to 200 ms steps from −135 mV to +60 mV in +15 mV increments applied every 5 s, from a holding voltage of either 0 mV or −50 mV. The absolute peak current from within the step was analyzed.

### Surface biotinylation

Three days after oocyte injections, 20–25 oocytes per sample were washed x6 in ND96 and labeled with 0.5 mg/ml EZ-Link™ Sulfo-NHS-SS-Biotin (Thermo Fisher) for 30 min at room temperature. The “no biotinylation” samples were incubated without the biotinylation reagent. Oocytes were washed again x6 in ND96 before homogenizing (by pipetting up and down 15 times with a P200 pipette) in 500 μL lysis buffer (1% Triton X-100, 100 mM NaCl, 20 mM Tris-HCl, pH 7.4) plus 1:1000 Protease Inhibitor Cocktail Set III, EDTA-Free (Calbiochem). All subsequent steps were performed at 4 °C. Lysates were gently shaken for 10 min then centrifuged at 16,000 x *g* for 5 min. A 50 μL aliquot of the supernatant (total cell protein) was stored at −80 °C for later use. The remaining supernatant was diluted 1:1 with the lysis buffer, then 100 μL of NeutrAvidin agarose beads (Thermo Fisher) added and the sample shaken gently overnight at 4 °C. For the desthiobiotin binding samples, 2.5 mM desthiobiotin was added at the bead incubation step. Beads were washed x6 in 500 μL lysis buffer with a 2 min 6,000 x *g* centrifugation between each wash, and finally resuspended in 50 μL lysis buffer. Both this sample and the total cell protein sample were mixed 1:1 with 2x loading buffer: 50% 4x Laemmli Sample Buffer (Bio-Rad), 30% 100 mM DTT, 10% 2-mercaptoethanol, and 10% Buffer BXT (IBA Lifesciences) and gently rotated for 1 h at room temperature. Samples were centrifuged for 2 min at 12,000 x *g* before being separated on 8–16% Mini-PROTEAN® TGX™ Precast Protein Gels (Bio-Rad) at 160 V for 60 min. PageRuler™ Plus Prestained Protein Ladder (Thermo Fisher) was used as the size standard. Samples were transferred onto a polyvinylidene difluoride membrane activated with methanol using the Trans-Blot Turbo Transfer System (Bio-Rad). Membranes were probed with mouse StrepMAB-Classic antibody diluted 1:3000 in TBS-T (25 mM Tris, 137 mM NaCl, 3 mM KCl, 0.05% Tween 20), followed by horseradish peroxidase conjugated anti-mouse secondary antibody diluted 1:10,000 in TBS-T, then developed using SuperSignal™ West Pico PLUS Chemiluminescent Substrate (Thermo Fisher).

### Baculovirus generation and expression of SARS-CoV-1 Orf3a_STREP_, SARS-CoV-2 Orf3a_STREP_

To produce high-titer SARS-CoV-1 Orf3a_STREP_, SARS-CoV-2 Orf3a_STREP_ and VPS39_GFP_ baculovirus, plasmids were first transformed into DH10Bac E. coli (Thermo Fisher) to generate bacmid DNA, using a blue-white colony selection strategy. Bacmid DNA was purified and transformed using CellFectin II reagent (Thermo Fisher) into SF9 cells resuspended in fresh ESF 921 Insect Cell Culture Medium, Protein Free (Expression Systems) and seeded in a 6-well plate to produce baculovirus. After 3-4 day incubation at 28°C (< 70% cell viability and abundant GFP expression), the supernatant containing a low titer of baculovirus (P0 virus) was collected and filtered. P0 virus was used to infect a larger suspension (100 mL) of SF9 cells to increase baculovirus titer (P1 virus), collected and filtered after 4 days of incubation at 28°C. A final round of baculovirus amplification in SF9 cells yielded 200-300 mL of baculovirus, supplemented with 5% FBS (P2 virus).

For protein expression, SARS-CoV-1 Orf3a or SARS-CoV-2 Orf3a P2 virus (3-10% of total cell suspension volume) was used infect Expi293F GnTI-cells, grown in FreeStyle 293 Expression Medium (Thermo Fisher) adapted to suspension growth in 2% Foundation FBS (GeminiBio). Cells were incubated in suspension for 24 hours at 37°C. Sodium butyrate (10 mM, Sigma-Aldrich) was added to enhance protein expression, and the cells were left for another 2-3 days incubating in suspension at 37°C. Cells were harvested by pelleting at 400 x *g*, flash-cooled and stored at −80°C. For cells overexpressing VPS39_GFP_, a 100 mL suspension was split into 10 mL aliquots and harvested as described.

### Purification of SARS CoV-1 and SARS-CoV-2 Orf3a

Cells (0.5-1L biomass) were agitated for 1 h at 4°C in Purification Buffer containing 100 mM Tris-HCl, pH 8.0, 75 mM NaCl, 75 mM KCl, 1 mM EDTA, pH 8.0, and supplemented with 25 µg/mL deoxyribonuclease I (DNase, Gold Biotechnology), a 1:1000 dilution of Protease Inhibitor Cocktail Set III, EDTA-free (PI Mix III, Calbiochem), 1 mM 4-(2-Aminoethyl)-benzenesulfonylfluoride hydrochloride (AEBSF, Gold Biotechnology), 1 mM benzamidine hydrochloride monohydrate (benzamidine, Gold Biotechnology), and 30 mM n-dodecyl-β-D-maltopyranoside (DDM, Anatrace). Following membrane protein extraction, the soluble fraction was clarified by centrifugation at 40,000 x *g* for 45 min at 4°C and vacuum filtered. For affinity purification with the twin-strep tag (22°C), lysate was sequentially applied over two columns packed with 1-2 mL of Strep-Tactin XT resin (IBA-Lifesciences), pre-equilibrated in Wash Buffer (Purification Buffer supplemented with 1 mM DDM). After application of lysate, the resin was rinsed with ∼5 column volumes of Wash Buffer and protein was eluted after a 30 min incubation of resin in 3 mL of Buffer BXT (IBA-Lifesciences) supplemented with 75 mM KCl and 1 mM DDM. The collected elution was concentrated to 6 mg/mL using a 50,000 Da concentrator (Amicon Ultra, EMD Millipore) and further purified by SEC using a Superdex 200 Increase 10/300 GL (Superdex 200, GE Healthcare) column in SEC buffer containing 20 mM HEPES, pH 7.5, 75 mM NaCl, 75 mM KCl, and supplemented with 3 mM n-decyl-β-D-maltopyranoside (DM, Anatrace).

### Expression and purification of MSP1D1

The nanodisc scaffold protein MSP1D1 was expressed and purified as described previously with modifications.^59^ MSP1D1 pET28a was transformed and expressed in *E. coli* B21-CodonPlus (DE3)-RIPL competent cells (Agilent). MSP1D1 contains a N-terminal 6xHis tag that can be removed by TEV protease. For purification, MSP1D1 *E. coli* pellets were agitated for 20 min at room temperature in buffer (25 mL/1 L of biomass) containing 20 mM sodium phosphate, pH 7.4 supplemented with 25 µg/mL DNase, a 1:1000 dilution of PI Mix III, 1 mM AEBSF, and 1 mM benzamidine. The resuspension was lysed by passing it 4x through a microfluidizer EmulsiFlex-C5 (Avestin) and clarified by centrifugation at 40,000 x *g* for 40 min at 4°C. For affinity purification, 300 mM NaCl was added to the lysate, combined with Ni-NTA resin (2 mL/1 L of biomass, Qiagen) pre-equilibrated with 40 mM sodium phosphate buffer, pH 7.4 and rotated for 1 h at 4°C. After batch binding and packing of the Ni-NTA resin, all subsequent wash and elution steps follow as described previously.^59^

To remove the 6xHis tag fused to MSP1D1, TEV protease was added to the elution fraction at a at a 1:20 wt:wt ratio, supplemented with 0.5 mM EDTA. The next day, TEV protease cleavage was confirmed by SDS-PAGE and the sample was dialyzed overnight in buffer containing 40 mM Tris-HCl, pH 8.0 and 300 mM NaCl. The following day, residual 6xHis-tagged TEV protease was removed by the addition of Ni-NTA resin, and the sample was incubated for 2 h at room temperature. After removing the resin, the sample was dialyzed overnight in storage buffer containing 20 mM HEPES, pH 7.4, 75 mM NaCl, 75 mM KCl and 0.2 mM DDM. The next day, the final sample was concentrated to ∼5 mg/mL in a 10,000 Da concentrator (Amicon Ultra, EMD Millipore), flash-cooled and stored at −80°C.

### Expression and purification of Saposin A

Saposin A was expressed and purified as described previously with modifications.^32, 33^ Saposin A pET28a was transformed and expressed in *E. coli* Rosetta-gami 2 (DE3) competent cells (Thermo Fisher). Saposin A contains a TEV protease-removable N-terminal 6xHis tag. For purification, MSP1D1 *E. coli* pellets were agitated for 20 min at 4°C in buffer (25 mL/1 L of biomass) containing 20 mM HEPES, pH 7.5, 150 mM NaCl, 20 mM Imidazole and 5% glycerol, supplemented with 25 µg/mL DNase, a 1:1000 dilution of PI Mix III, 1 mM AEBSF, and 1 mM benzamidine. The resuspension was lysed by passing it 4x through a microfluidizer and first clarified by centrifugation at 2000 x *g* for 30 min at 18°C to remove insoluble cell debris. The sample was heated at 85°C for 10 min, followed by a second clarification step by centrifugation at 55,000 x *g* for 45 min at 18°C. For affinity purification, supernatant was combined with Ni-NTA resin (1 mL/1 L of biomass, Qiagen) pre-equilibrated with 20 mM HEPES, pH 7.5 and 150 mM NaCl and rotated for 1 h at 4°C. After batch binding and packing of the Ni-NTA resin, all subsequent wash and elution steps follow as described previously.^32^

To remove the 6xHis tag fused to Saposin A, TEV protease was added to the elution fraction at a 1:10 wt:wt ratio. The sample was dialyzed overnight in a 3.5kDa molecular weight cutoff dialysis cassette (Slide-a-Lyzer, Thermo Fisher) at 4°C against 20 mM HEPES, pH 7.5 and 150 mM NaCl. The next day, the sample was passed over a Ni-NTA column to remove uncleaved Saposin A and 6xHis-tagged TEV protease, followed by SDS-PAGE to confirm cleavage. This sample was dialyzed in the same type of dialysis cassette overnight in storage buffer containing 20 mM HEPES, pH 7.5, 75 mM NaCl and 75 mM KCl. The next day, 0.2 mM DDM was added prior to concentration to prevent aggregation. The final sample was concentrated to ∼2.8 mg/mL (10,000 Da concentrator, Amicon Ultra, EMD Millipore), flash-cooled and stored at −80°C.

### Orf3a reconstitution into MSP1D1 and Saposin A nanodiscs

SARS-CoV-1 Orf3a and SARS-CoV-2 Orf3a were reconstituted into MSP1D1 nanodiscs containing two distinct lipid compositions which we refer to as ‘plasma membrane’ (PM) or ‘late endosomal/lysosomal’ (LE/Lyso). A 2:1:1 (wt:wt) PM lipid mixture of 1,2-dipalmitoleoyl-sn-glycero-3-phosphocholine, 1,2-dioleoyl-sn-glycero-3-phosphoethanolamine, and 1,2-dioleoyl-sn-glycero-3-phospho-L-serine (16:1 PC:DOPE:DOPS, Avanti Polar Lipids) or a 4:2:1 (wt:wt) LE/lysosome lipid mixture of 16:1 PC, DOPE, and (S,S) bisoleoyl-lysobisphosphatidic acid (LBPA, Echelon Biosciences) was prepared at 10 mM in SEC buffer with 5 mM DM. For MSP1D1 nanodiscs, lipid nanodiscs were formed by first combining Orf3a, MSP1D1 and the lipid mixture together at a molar ratio of 1:4:260 and rotating at 4°C for 1 h. For Saposin A nanodiscs, a molar ratio of 1:15:40 of Orf3a: Saposin A: lipid was used to form protein-reconstituted nanodiscs. First, SARS-CoV-2 Orf3a was added to the lipid mixture and incubated at room temperature for 10 min. Saposin A was then added to the mixture and incubated for 30 min at room temperature. For both MSP1D1 and Saposin A nanodiscs, detergent was removed to form lipid nanodiscs by the addition of bio-beads (Bio-beads SM2 resin, Bio-rad) pre-equilibrated in SEC buffer (∼1:3, wt:vol). Bio-beads were added and incubated with the sample rotating at 4°C for 1 h. Then, a second round of Bio-Bead binding was performed, and the sample was left rotating at 4°C overnight. The next day, the lipid nanodiscs were purified by SEC in detergent-free SEC buffer. The peak fraction was collected in each case, concentrated to 1.3-1.5 mg/mL using a 100,000 Da concentrator (Amicon Ultra, EMD Millipore), and immediately used for cryo-EM grid preparation.

### EM sample preparation and data acquisition

3 µL of sample was pipetted onto Quantifoil R1.2/R1.3 holey carbon grids (Cu 400, Electron Microscopy Sciences) which had been glow discharged for 1 min using a PELCO easiGlow glow discharge cleaning system (Ted Pella). A vitrobot Mark IV cryo-EM sample plunger (FEI) (operated at 4°C with a 4-5 s blotting time at a blot force of 3-4 and 100% humidity) was used to plunge-freeze the sample into liquid nitrogen-cooled liquid ethane. Grids were first imaged on a 200 keV Technai F20 TEM (FEI) equipped with a K2 Summit direct electron detector (Gatan, Inc.) to evaluate sample quality, particle quantity and distribution. They were subsequently clipped and loaded into a 300 keV Titan Krios microscope (FEI) equipped a Cs corrector to reduce spherical aberration of the objective lens from 2.7 mm to ∼0.01 mm, BioQuantum energy filter (Gatan, Inc.) and a K3 camera (Gatan, Inc.). Images were recorded with SerialEM in super-resolution mode at a calibrated magnification of 59,242x, which corresponds to a super-resolution pixel size of 0.422 Å, a defocus range of −0.8 to −2 µm, and a total e^-^ dose of 50 e^-^ per Å^2^.^60^

### Cryo-EM data processing

*Figure 3-figure supplement 1, 3* and *Figure 4-figure supplement 2*, 4 describe the workflow for SARS-CoV-2 Orf3a LE/Lyso MSP1D1 nanodisc, SARS-CoV-2 Orf3a PM MSP1D1 nanodisc, SARS-CoV-2 Orf3a LE/Lyso Saposin A nanodisc and SARS-CoV-1 Orf3a LE/Lyso MSP1D1 nanodisc datasets. Movie stacks were gain-corrected, two-fold Fourier-cropped to a calibrated pixel size of 0.844 Å/pix and dose weighted using the CPU-based Relion 3.0 (SARS-CoV-2 Orf3a LE/Lyso MSP1D1 nanodisc) or Relion 3.1 (SARS-CoV-2 Orf3a PM MSP1D1 nanodisc, SARS-CoV-2 Orf3a LE/Lyso SaposinA nanodisc and SARS-CoV Orf3a LE/Lyso MSP1D1 nanodisc) implementations of MotionCor2.^61, 62^ Contrast Transfer Function (CTF) estimates for motion-corrected micrographs were performed in CTFFIND4 using all 50 frames.^63^

### SARS-CoV-2 Orf3a LE/Lyso MSP1D1 nanodisc dataset

Two datasets of the SARS-CoV-2 Orf3a LE/Lyso nanodisc preparation (SARS-CoV-2 Orf3a LE/Lyso datasets A and B) were collected with a total of 8442 and 5645 micrographs, respectively. Micrographs with CTF estimation fits worse than 5 Å or with a prominent ice ring as manually inspected from the CTF 2D power spectra were discarded, leaving a total of 6759 and 3033 good micrographs per dataset.

Initial data processing was performed with SARS-CoV-2 Orf3a LE/Lyso dataset A. To assess the oligomeric state of SARS-CoV-2 Orf3a, all steps during the initial cycle of data processing were performed without symmetry applied. 1123 particles were manually selected for reference-free 2D classification to generate templates that were used for automatic particle picking with a subset (∼300) of micrographs in Relion 3.0.^61^ Approximately 100,000 particles were selected and were used to improve the templates generated by reference-free 2D classification. The improved templates were used for automatic particle picking with the entire dataset. 2,520,325 particles were subjected to two rounds of reference-free 2D classification in cryoSPARC3.0 to remove outlier particles (e.g. remove ice contaminants), yielding 1,042,957 particles.^64^ An initial model of SARS-CoV-2 Orf3a was generated by *ab initio* reconstruction in cryoSPARC3.0.^64^ Three rounds were performed using low- and high-resolution cutoffs of 9 Å and 7 Å, initial and final particle batch sizes of 1000 and 2000 particles, and a class similarity of 0.01 for each iteration. Symmetry was not applied at this stage. Using non-uniform refinement in cryoSPARC3.0, the best initial 3D model from ∼86,126 particles refined to ∼6 Å and showed clear two-fold symmetry.^65^

To further improve the model, additional particles from the second round of reference free 2D classification were selected for (1,363,667 particles in total) and subjected to another round of 2D classification in cryoSPARC3.0. A similar strategy of 4 rounds of by *ab initio* reconstruction was performed, with the final iteration using an initial and final particle batch size of 2000 and 4000 particles, and a class similarity of 0.4. The best *ab initio* model from the last iteration was refined to 5.3 Å from 73,377 particles using non-uniform refinement with C2 symmetry applied. The particles were then subjected to Bayesian polishing in Relion 3.0 and refined in cryoSPARC3.0, to yield a final reconstruction of similar resolution.^61^

A data processing ‘decoy’ refinement strategy was implemented at this step to improve map resolution as described previously.^66^ Micrographs were over-picked with particles using a Laplacian of Gaussian filter in Relion 3.0.^61^ Particles were subjected to a round of reference-free 2D classification in cryoSPARC3.0 to remove ice contaminants. The starting particle stack of 6,145,296 was used to generate seven junk ‘decoy’ models from a cycle of *Ab initio* reconstruction. The decoy maps and best SARS-CoV-2 Orf3a map (∼5.3 Å) were used to sort false positive from Orf3a particles using heterogeneous refinement in cryoSPARC3.0 (Round 1).^64^ Orf3a particles were subjected to additional rounds of heterogeneous refinement until most particles classified with the SARS-CoV-2 Orf3a model, with the final particle stack containing 1,565,551. Iterative rounds of *ab initio* reconstruction were performed to further classify this subset of particles using similar parameters to what was described above. The best 3D *ab initio* model from 300,114 particles was refined to 4.1 Å using non-uniform refinement in cryoSPARC3.0. Particles that had an inter-particle distance of <100 Å (particle duplicates) were removed and subjected to Bayesian polishing and 3D refinement in Relion 3.0. The final 3D reconstruction resolved to 3.8 Å.

Another iteration (Round 2) of the ‘decoy’ refinement strategy was performed with the starting particle stack. Three rounds of heterogeneous refinement using the improved SARS-CoV-2 Orf3a model at 3.7 Å were completed, followed by an additional reference-free 2D classification in cryoSPARC3.0 with the 1,184,876 particles to remove outliers.^64^ The selected particles from 2D classification (1,049,046 particles – eventually combined with dataset B) yielded a refined 3D reconstruction of ∼4 Å. Particle duplicates were removed and the remaining particles were subjected to Bayesian polishing and a round of 3D refinement in Relion 3.0. To assess distinct conformational states, we performed 3D classification in Relion 3.0 using the consensus reconstruction as an initial model (low-pass filtered at 5 Å resolution) and applied an Orf3a-only mask. The particles were sorted into 3-4 classes using the fixed angular assignments from 3D refinement. Although this strategy did not yield a different conformation of Orf3a, it did help to identify a set of 186,327 particles that generated a reconstruction which refined to 3.8 Å with more 3D features.

At this stage, CoV2 Orf3a LE/Lyso dataset B was combined with dataset A. Particles were automatically selected in dataset B using the Laplacian of Gaussian filter strategy in Relion 3.0. The dataset B particles were then subjected to the ‘decoy’ refinement strategy using the improved model of Orf3a (3.8 Å). The particles that were classified with the SARS-CoV-2 Orf3a model (962,666 in total) were combined with the 1,049,046 particles from dataset A, refined to ∼4 Å resolution. These were processed with Bayesian polishing in Relion 3.0 and imported to cryoSPARC3.0 for sorting by 2D classification and *Ab initio* reconstruction. The best model reconstructed from 520,044 particles using non-uniform 3D refinement reached a resolution of 3.5 Å. The parameters from Bayesian polishing were recalculated for this subset of particles, and the particles were subjected a round of 3D refinement and 3D classification with the regularization parameter T adjusted to a value of 20 to sort out the best model at 3.4 Å resolution reconstructed from 119,318 particles.

A final iteration of data processing was performed using Topaz, a neural network particle picking program implemented in cryoSPARC3.0.^67^ Topaz denoise was first used to reduce noise in all micrographs prior to particle picking.^68^ The best 119,318 particles from the last round of data processing were used to train the ResNet8 neural network to select particles from CoV2 Orf3a LE/Lyso datasets A and B.^67^ The 7,557,588 particles selected were subjected to reference-free 2D classification to first remove ice contamination, followed by a ‘decoy’ refinement strategy using the best 3.4 Å resolution model. The particles that sorted with the SARS-CoV-2 Orf3a model (1,430,051 in total) were further sorted by one round of *ab initio* reconstruction to identify the best subset of particles. The final reconstruction from a total of 679,097 particles in the best *ab initio* class refined to 3.4 Å resolution. Particle duplicates were removed, and the remaining particles were subjected to Bayesian polishing and a round of 3D refinement in Relion 3.0. This was followed by 3D classification in searching for 3 classes, with the regularization parameter T adjusted to a value of 40. The best map and particles from 3D classification were used for 3D refinement, CTF refinement, and a final round of 3D refinement in Relion 3.0 to yield a final map of 3.0 Å resolution.

### SARS-CoV-2 Orf3a PM MSP1D1 nanodisc dataset

A dataset of the SARS-CoV-2 Orf3a PM nanodisc preparation was collected with a total of 15,637 good quality micrographs. Micrographs with CTF estimation fits less than 5 Å, a rlnCtfAstimatism value greater than 1000 Å, rlnFigureofMerit values below 0.1, or with a prominent ice ring as manually inspected from the CTF 2D power spectra, were discarded.^57^ Particle picking was performed using Topaz implemented in cryoSPARC3.0.^67^ The micrographs were first denoised and 1678 particles were manually selected to train the ResNet8 neural network for particle selection.^67, 68^ The 8,607,878 particles selected were subjected to reference-free 2D classification to first remove ice contamination, and were then sorted through iterative rounds of heterogeneous refinement with 7 ‘decoy’ maps and the a 3.6 Å resolution map of the SARS-CoV-2 Orf3a LE/Lyso MSP1D1 nanodisc.^60^ An initial model from the particles which sorted into the “good class” (475,773 particles) was generated by *ab initio* reconstruction and refined to 3.8 Å resolution using non-uniform refinement in cryoSPARC.^60^ The initial model was mildly improved in Relion 3.1, through Bayesian polishing, 3D refinement and particle sorting by 3D classification, searching for 3 classes with fixed angular distribution and T=40 parameters.^57^ A final reconstruction of 3.7 Å from 129,484 particles was used as inputs for the next round of particle picking and heterogeneous refinement.

The 129,484 particles from the first iteration Topaz particle picking and map improvement were used to train the ResNet8 neural network in Topaz. A total of 13,224,923 particles were selected and sorted by 2D classification and iterative rounds of 3D heterogeneous refinement with the decoy and SARS-CoV-2 Orf3a PM MSP1D1 nanodisc 3.7 Å maps. A total of 595,295 particles were selected and used for non-uniform in refinement in cryoSPARC3.0, which generated a 3.7 Å reconstruction.^65^ This was improved in in Relion 3.1 with Bayesian polishing, 3D refinement, two rounds of 3D classification in searching for 4 classes each time by fixing the angular distribution and T=40 parameters. The final model, reconstructed from 125,678 particles, was subjected to 3D refinement, CTF refinement, another round of Bayesian polishing, and 3D refinement to yield a final reconstruction of 3.4 Å resolution.

### SARS-CoV-2 Orf3a LE/Lyso Saposin A nanodisc dataset

Two datasets of the SARS-CoV-2 Orf3a Saposin A nanodisc preparation was collected with a total of 15946 good quality micrographs. Micrographs with CTF estimation fits worse than 5 Å, a rlnCtfAstimatism value greater than 1000 Å, a rlnFigureofMerit value that was below 0.1, or with a prominent ice ring as manually inspected from the CTF 2D power spectra were discarded.^57^ Particle picking was performed using Topaz.^67^ The micrographs were first denoised, and 1297 particles were manually selected to train the ResNet8 neural network to select particles.^67, 68^ The 12,070,379 particles selected were subjected to reference-free 2D classification to first remove ice contamination.^60^ An initial model of a SARS-CoV-2 Orf3a LE/Lyso Saposin A nanodisc was generated, first by *ab initio* reconstruction using cryoSPARC3.0, using the same parameters described above and 8,303,729 particles.^60^ This was followed by non-uniform refinement in cryoSPARC3.0 with the best 3D map from *ab initio* reconstruction. 1,907,669 particles refined to ∼3.1 Å and showed clear low-resolution density for 6 Saposin A molecules.^65^

This model was used with 7 decoy models to further parse out the best particles from a 11,247,076 stack using iterative heterogeneous refinement.^60^ The best particles were subjected to 2D classification to further remove ice, another round of *ab initio* reconstruction, and the best map and particles from the best 3 classes generated a reconstruction to ∼ 2.9 Å resolution containing 2,559,504 particles. To improve the map and look for other conformational states of SARS-CoV-2 Orf3a we performed subsequent data processing steps in Relion 3.1.^57^ Particles duplicates were removed and then subjected to Bayesian polishing and 3D refinement. Next, two individual rounds of 3D classification were performed, searching for 6 or 10 classes with fixed angular distribution and T=40 parameters and an initial low resolution pass filter of 8 Å or 5 Å, respectively. We were unable to identify any alternative conformations. The best map resolved to 2.8 Å resolution after CTF refinement, 3D refinement, another round of Bayesian polishing and a final round of 3D refinement.

### SARS-CoV-1 Orf3a LE/Lyso nanodisc datasets

A dataset of the SARS-CoV-1 Orf3a LE/Lyso nanodisc preparation was collected with a total of 13,970 good quality micrographs. Micrographs with CTF estimation fits >5 Å, a rlnCtfAstimatism value greater than 1000 Å, rlnFigureofMerit value that was below 0.1, or with a prominent ice ring as manually inspected from the CTF 2D power spectra were discarded.

Particle picking was performed using Topaz.^67^ The micrographs were first denoise, and 1504 particles were manually selected to train the ResNet8 neural network to select particles.^67, 68^ The 4,697,964 particles selected were subjected to reference-free 2D classification to first remove ice contamination.^60^ An initial model of a SARS-CoV-1 Orf3a LE/Lyso MSP1D1 nanodisc was generated, first by two rounds of *ab initio* reconstruction using cryoSPARC3.0 with 2,702,734 particles and 5-6 classes. ^60^ This was followed by 2D classification to further remove ice, and another round of *ab initio* reconstruction searching for 3 classes with 620,611 particles. The best map reconstructed from 156,694 particles was refined to ∼4.5 Å using non-uniform refinement, and then further improved to ∼3.6 Å by Bayesian polishing and 3D refinement in Relion 3.1.^61, 65^

The final map generated from iterative *ab initio* reconstruction was then used with 7 decoy maps for sorting 3,908,602 particles by iterative heterogeneous refinement.^60^ The best particles from this procedure were refined, particle duplicates were removed, and the map was improved by Bayesian polishing, 3D refinement and 3D classification, searching for 3 classes with fixing the angular distribution and T=40 parameters in Relion3.1.^61^ A subset of 15,848 particles was identified that generated a 3.6 Å reconstruction after 3D refinement. This stack of particles and map was then used to train the Topaz ResNet8 neural network to re-select particles, and for iterative heterogeneous refinement. This entire procedure was repeated twice more and the final map, which was selected from a 3D classification step in Relion3.1, followed by CTF refinement, 3D refinement, Bayesian polishing and final 3D refinement to generate a reconstruction to 3.1 Å with 162,607 particles.

### Model building and refinement

The atomic models were manually built into one of the half-maps that had been ‘postprocessed’ in Relion3.0 or Relion 3.1 using a *B*-factor of −50 Å^2^ and low-pass filtered at the final overall resolution, and the same half map that had been density-modified using resolve-cryo-EM in PHENIX.^61, 69^ The high-resolution structure of SARS CoV-2 Orf3a was used as a starting point (PDB ID: 7KJR) and refined in real space using the COOT software.^70^ The atomic models were further refined in real space against the same half-map using real-space-refine-1.19 in PHENIX.^71^ The final models have good stereochemistry and good Fourier shell correlation with the other half-map as well as the combined map (*Figure 3-figure supplement 2, Figure 4-figure supplement 3*). Structural figures were prepared with Pymol (pymol.org), Chimera, HOLE and APBS.^72–74^

### Orf3a reconstitution into liposomes for functional assays

SEC-purified protein [in SEC buffer containing 3 mM DM (Anatrace)] was reconstituted into liposomes. For flux assays, a 3:1 (wt/wt) mixture of 1-palmitoyl-2-oleoyl-sn-glycero-3-phosphocholine (POPE, Avanti) and 1-palmitoyl-2-oleoyl-sn-glycero-3-phospho-(1’-rac-glycerol (POPG, Avanti) lipids) was prepared at 20 mg/mL in reconstitution buffer (see flux assay protocols below). 8% (wt/vol) n-octyl-b-D-maltopyranoside (OM, Anatrace) was added to solubilize the lipids, and the mixture was incubated with rotation for 30 min at room temperature. Purified protein was mixed with an equal volume of the solubilized lipids to give a final protein concentration of ∼0.08-0.8 mg/mL and a lipid concentration of 10 mg/mL (at ∼1:100 or 1:10, wt:vol ratio). Proteoliposomes were formed by dialysis (using an 8000 Da molecular mass cutoff, Spectrum Labs) for 4-5 days at 4 °C against 4 L of reconstitution buffer and flash-cooled and stored at −80 °C until use. For the flux assay experiment that included LBPA, a 6:2:1 mixture of POPE, POPG and LBPA was prepared, and this mixture was used for Orf3a reconstitution. For all experiments, a liposome-only control was prepared concurrently in the absence of protein.

For proteoliposome patch clamp, soybean polar lipid extract (Asoletin, Avanti Polar Lipids) was prepared at 10 mg/mL in reconstitution buffer including 200 mM KCl, 10 mM HEPES, pH 7.4. Purified protein was concentrated to ∼0.08-0.8 mg/mL and mixed dropwise with an equal volume of asolectin (at ∼1:100 or 1:10, wt:vol ratio), diluted ∼5-10 fold in reconstitution buffer, and transferred to a dialysis cassette (10,000 Da molecular mass cutoff Slide-A-Lyzer, Thermo-Fisher). Proteoliposomes were formed by detergent dialysis for 4-5 days at 4 °C against 4 L of reconstitution buffer. After dialysis, proteoliposomes were concentrated to a final lipid concentration of 50 mg/mL by ultracentrifugation at 100,000 x *g* for 1 h at 4°C, followed by resuspension and sonication in the appropriate volume of fresh reconstitution buffer. Samples were aliquoted, flash-cooled and stored at −80°C. For all experiments, a liposome-only control was prepared concurrently in the absence of protein.

### ACMA fluorescence-based flux assay

The ACMA fluorescence-based flux assays were performed as described previously with modification.^75–78^ Briefly, Orf3a was reconstituted into liposomes by dialysis in buffer containing 150 mM KCl and 10 mM HEPES, pH 7.0 (pH-adjusted with N-methyl-D-glucamine, NMDG) to evaluate K^+^ flux or 100 mM Na_2_SO_4_ and 10 mM HEPES, pH 7.0 (pH-adjusted with NaOH) to test for Cl^-^ flux. Reconstituted Orf3a was quickly thawed, sonicated, and diluted 1:100 into Flux Assay Buffer (FAB) containing 150 mM NMDG-Cl, 10 mM HEPES, pH 7.0, 0.5mg/mL BSA, and 2 µM 9-Amino-6-Chloro-2-Methoxyacridine (ACMA, Sigma-Aldrich) to establish a K^+^ gradient. A similar FAB was prepared to create a Cl^-^ gradient by substituting 150 mM NMDG-Cl with 125 mM NaCl. The assay was performed in a 96-well format using a CLARIOstar Plus microplate reader (BMG Labtech) with excitation and emission wavelengths set to 410 nm and 490 nm, respectively. Fluorescence was recoded in kinetics mode for 20 min with 30 s between reach read. After 150 s, the proton ionophore carbonyl cyanide m-chlorophenyl hydrazone (CCCP, Sigma-Aldrich) was added at 1 µM to each sample to promote the uptake of protons (H^+^) due to the electrochemical gradient established by efflux (K^+^) or influx (Cl^-^) of ions through Orf3a. At the end of the K^+^ flux assay (930 s), 20 nM valinomycin (Sigma-Aldrich) was added to each sample to release all K^+^ from vesicles.

### 90° light-scattering flux assay

The 90° light-scattering flux assay was adapted from a published protocol.^36, 37^ Orf3a was reconstituted in dialysis buffer containing 200 mM K-glutamate and 20 mM HEPES, pH 7.2. Liposomes were thawed, sonicated, and diluted 1:200 into a hypertonic FAB containing 260 mM K-thiocyanate (KSCN) and 20 mM HEPES, pH 7.2, in a stirred cuvette. 90° scattered light was recorded for 10 min using a Varian Carey UV-Vis Spectrophotometer (Agilent) with excitation and emission wavelengths at 600 nm. After 480 s, 20 nM valinomycin (Sigma-Aldrich) was added to each sample to measure K^+^ influx for all vesicles in the sample, resulting in vesicle swelling and a reduction in light scattering.

### Proteoliposome patch clamp experiments

Excised proteoliposome voltage clamp recordings were performed similar to protocols previously described.^3,79^ Briefly, purified SARS-CoV-2 Orf3a was reconstituted in soybean polar lipid extract (Avanti Polar Lipids) at 1:10 and 1:100 wt:wt protein to lipid ratios. Samples of this reconstitution were then dried on a cover glass under continuous vacuum at 4°C for 12 h. The dried aliquots were then rehydrated for 40-50 min with rehydration buffer: 200 mM KCl, and 5 mM HEPES at pH 7.2. A suspension of the rehydrated proteoliposomes was collected and used to seed a recording chamber cover glass. Once the proteoliposomes settled (15-20 minutes), multi-giga-ohm seals were achieved with SylGard coated pipettes pulled and polished to 2-4 megohms. The excised patch configuration was then achieved by retracting the pipette and breaking the connecting membrane with a physical jolt to the head stage. For symmetric recording conditions, internal and external solutions were as follows: Sodium solution; 150 mM NaCl, 1 mm MgCl_2_, 1 mM CaCl_2_, and 5 mM HEPES: Potassium solution; 150 mM KCl, 1 mm MgCl_2_, 1 mM CaCl_2_, and 5 mM HEPES: Calcium solution; 100 mM CaCl_2_, 1 mM MgCl_2_, and 5 mM HEPES. Voltage-clamp recordings were performed using an Axopatch 200B in patch configuration and the data acquired via Digidata 1440A and Clampex 10.7 software (Molecular Devices). High resolution current-voltage step protocols were performed at 125 kHz sampling rate with a 5 kHz Bessel Filter. Extended/time-course recordings were acquired using 1 s repeated step protocols with a 10 kHz sampling rate and 2 kHz Bessel filtering. Gap-free recordings were performed at 100 kHz with a 5 kHz Bessel filter. Analysis of single channel sweeps was performed using ClampSeg GUI idealization, as well as custom manual thresholding and delta-search scripts, to calculate the conductance and dwell time of all single channel events present.^80, 81^ Event probabilities were then calculated as the mean open probability of a given conductance across all successful excised patch recordings.

### Mass Spectrometry

Mass spectrometry and data analysis of SARS-CoV-2 Orf3a vesicles reconstituted at a high protein to lipid ratio were performed by the Proteomics Core Facility at the Whitehead Institute (Boston, MA).

### Co-immunoprecipitation experiments

Each aliquot of cells expressing VPS39_GFP_ were resuspended in 5 mL of buffer containing 50 mM HEPES, pH 7.5, 75 mM NaCl, 75 mM KCl, 1 mM EDTA, pH 8.0, a 1:1000 dilution of PI Mix III and 0.5% NP-40 (Millipore Sigma). Samples were incubated on dry ice for 10-20 min to lyse and then divided into 800 μL aliquots. For binding with SARS-CoV-1 or SARS-CoV-2 Orf3a constructs, protein purified in DM (0.5-50 μg), or a DM-buffer control, was added to each sample and rotated for 1 h, 4°C. Each sample was centrifuged at 14,000 x *g* for 15 min to remove cell debris. Input samples for the co-IP were taken for examination on a gel at this stage. The rest of the sample was incubated with 50 μL bed volume of StrepTactinXT for 2 h, 4°C. The flow-through after binding was collected at this stage. Resin was washed 5x with wash buffer containing 50 mM HEPES, pH 7.5, 75 mM NaCl, 75 mM KCl, 1 mM EDTA, pH 8.0, and 0.05% NP-40. The third wash step was used as a wash control. Wash buffer was added, mixed 1:1 with 2x loading buffer (50% Laemmli Sample Buffer, 500 mM DTT) to the resin and collected.

The western blot protocol is similar to that described for the surface biotinylation experiments. Samples were separated on 4–20% Mini-PROTEAN® TGX™ Precast Protein Gels 150 V for 60 min. For streptavidin detection, membranes were probed with mouse StrepMAB-Classic antibody diluted 1:5000 in TBS-T, followed by horseradish peroxidase conjugated to anti-mouse secondary antibody and developed using SuperSignal™ West Pico PLUS Chemiluminescent Substrate. For GFP detection, membranes were probed with rabbit monoclonal GFP antibody diluted 1:4000 in TBS-T, followed by Horseradish Peroxidase conjugated anti-rabbit secondary antibody at 1:10000 dilution in TBS-T and developed using SuperSignal™ West Pico PLUS Chemiluminescent Substrate.

## Acknowledgements

We’d like to thank Chris Miller for consultation on the ion channel reconstitution and the 90° light-scattering flux assay and Luke Lavis for use of their Varian Carey UV-Vis Spectrophotometer for these experiments. We also thank Lukas Frey for helpful suggestions with MSP1D1 nanodisc preparations, Kathy Schaefer for providing her expertise in FACS sorting, Eric Spooner for assistance with mass spectrometry, and William Patton of Scientific Computing at the Janelia Research Campus for assistance with single-channel recording analysis. We would also like to acknowledge cryoEM facility staff members Momoko Shiozaki and Shixin Yang for providing technical and computational support. Finally, we would like to thank members of the Clapham Lab for productive discussion, and in particular Ray Hulse for insight into structural interpretation and Shu-Hsien Sheu for helpful suggestions with collection and processing of confocal imaging data. B.C.A. received funding from the Australian National Health and Medical Research Council CJ Martin Fellowship (APP1162427). S.E.B. received funding from The Millennium Nucleus of Ion Channel-Associated Diseases (MiNICAD, #NCN19_168), a Millennium Nucleus of the Iniciativa Milenio, National Agency of Research and Development (ANID, Chile).

## Competing interests

The authors do not declare any competing interests.

---

**Figure 1-figure supplement 1.**
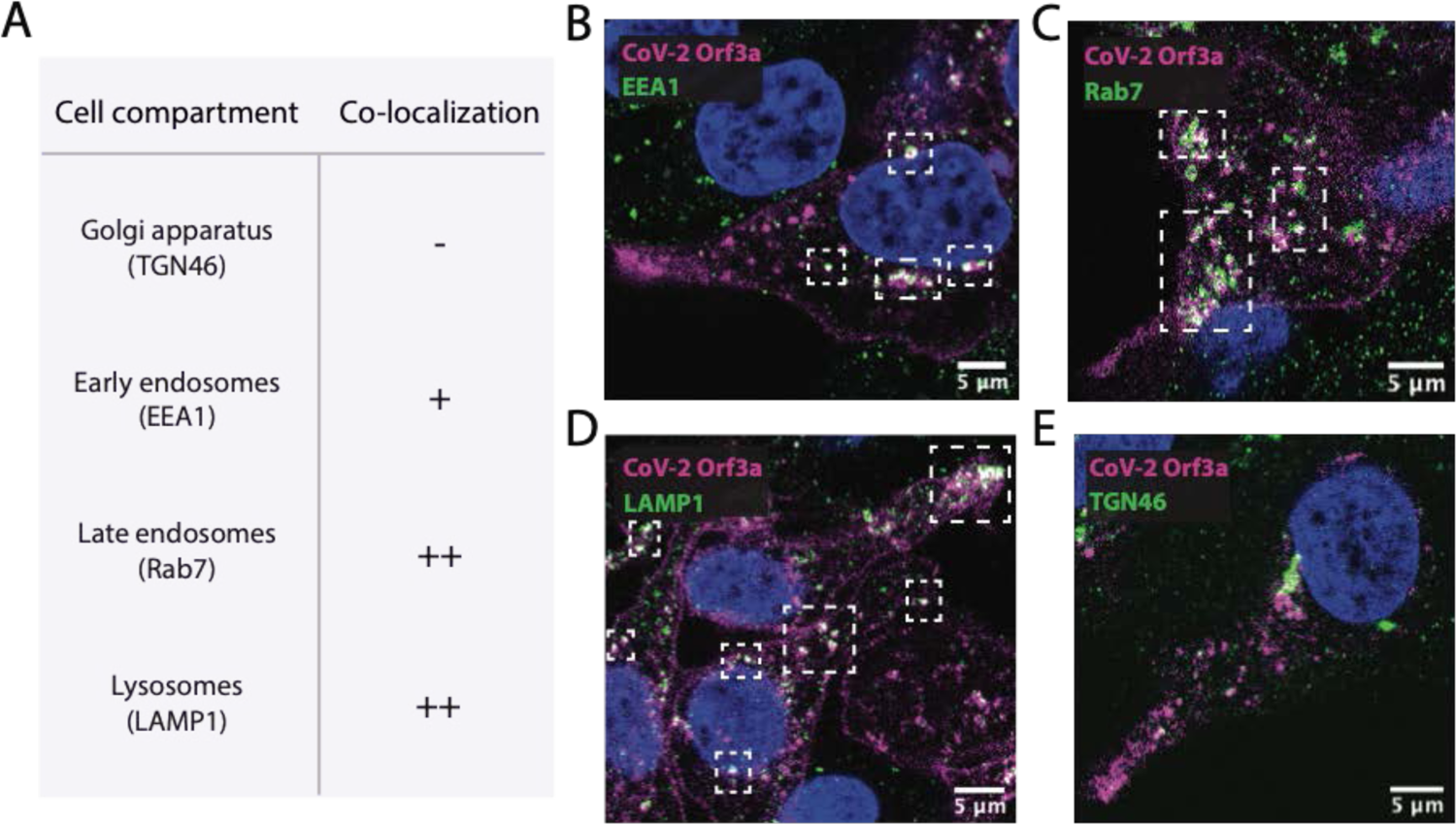
Sub-cellular localization by immunostaining of SARS-CoV-2 Orf3a (**A**) SARS-CoV-2 (CoV-2) Orf3a_HALO_ colocalizes with markers of the endocytic pathway, with minor overlap with a marker for Golgi. All primary antibody markers used to identify cellular compartments are listed in the table in A. (**B-E**) Fixed HEK293 cells treated with doxycycline to induce expression of CoV-2 Orf3a_HALO_ (CoV-2 3a, magenta) stained with (**B**) EEA1, (**C**) Rab7, (D) LAMP1, and (**E**) TGN46 (green). Nuclei are labeled with Hoechst 33258 (blue).

**Figure 1-figure supplement 2.**
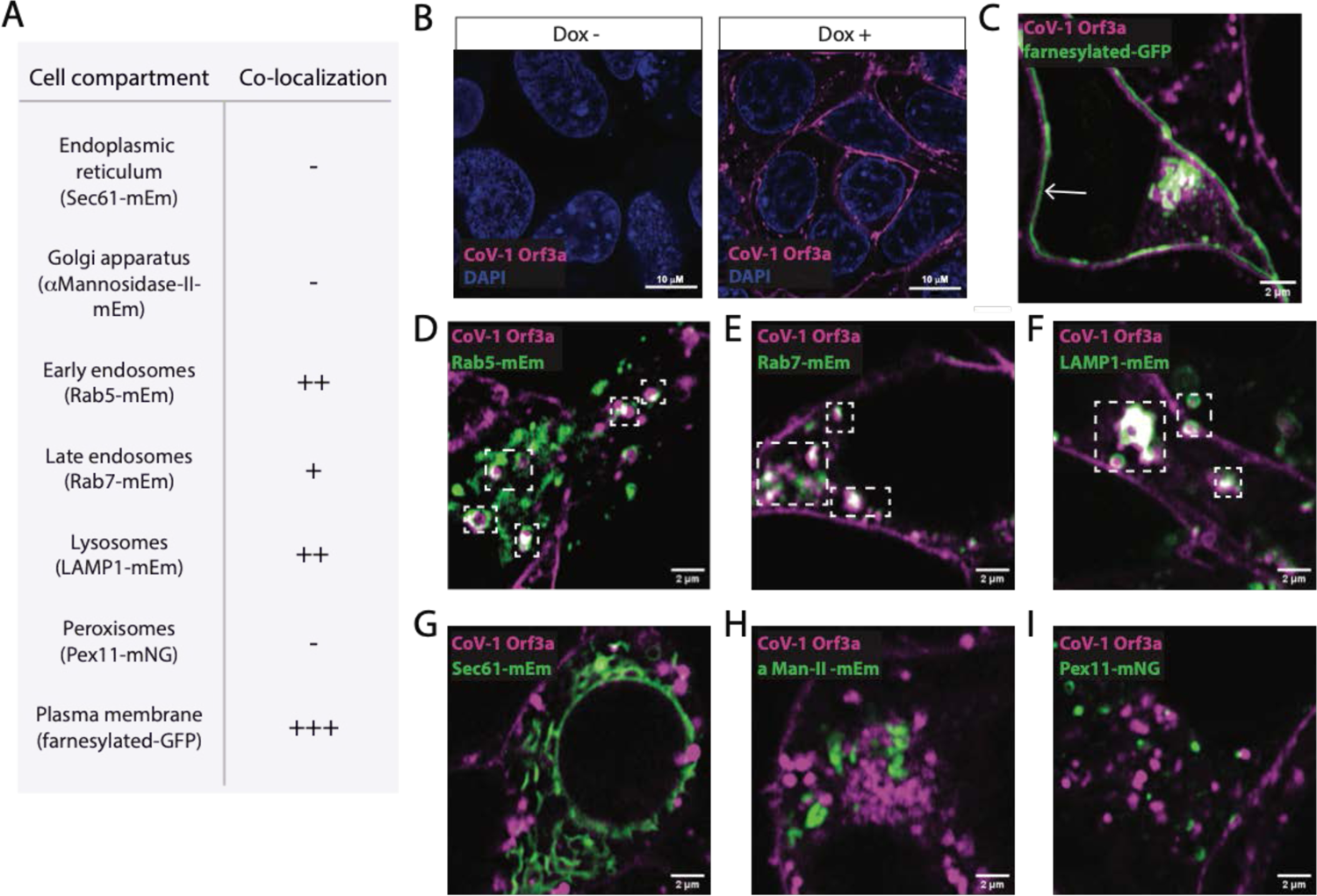
Sub-cellular localization of SARS-CoV-1 Orf3a (**A**) SARS-CoV-1 (CoV-1) Orf3a colocalizes with markers for the PM and endocytic pathway. All proteins markers used to identify cellular compartments are listed in the table (**A**) and are transiently expressed (mEm, mEmerald; mNG, mNeonGreen) (**B**) Live cell imaging of HEK293 cells without (Dox -) and with (Dox +) doxycycline to induce expression of CoV-1 Orf3a_HALO_ (magenta). Cell nuclei are visualized with Hoechst 33258 stain. (**C-I**) Live-cell image of transiently expressed (**C**) farnesylated-GFP, (**D**) Rab5-mEmerald, (**E**) Rab7-mEmerald, (**F**) LAMP1-mEmerald, (**G**) Sec61-mEmerald (**H**) aMannosidase-II-mEmerald or (**I**) Pex11-mNeonGreen (green) with CoV-1 Orf3a_HALO_ (magenta) using the HEK293 stable cell line in (**B**). White arrows and boxes indicate regions of co-localization.

**Figure 2-figure supplement 1.**
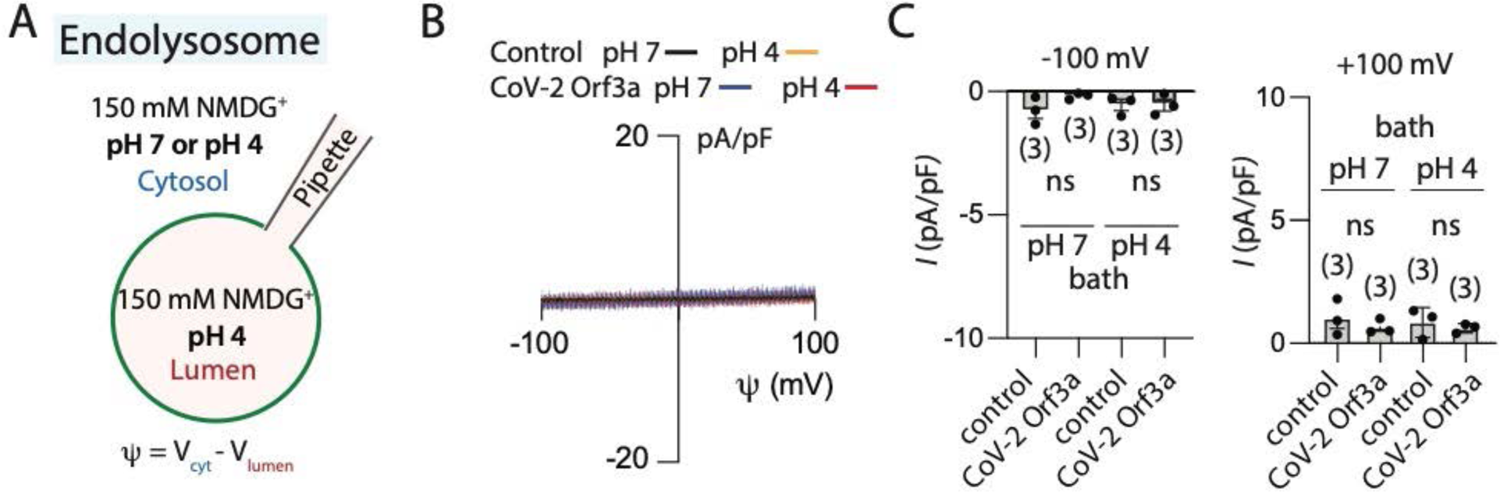
SARS-CoV-2 Orf3a_HALO_ does not elicit an H^+^-selective current in endo-lysosomes. (**A**) Internal and external recording solutions used in the endolysosomal patch clamp experiment. (**B**) I-V relationship for untransfected HEK293 cells (control, black) and transiently expressing SARS-CoV-2 (CoV-2) Orf3a_HALO_. (**C**) Average current density for untransfected HEK293 cells (control) and CoV-2 Orf3a_HALO_ (Orf3a) at −100 mV and +100 mV.

**Figure 2-figure supplement 2.**
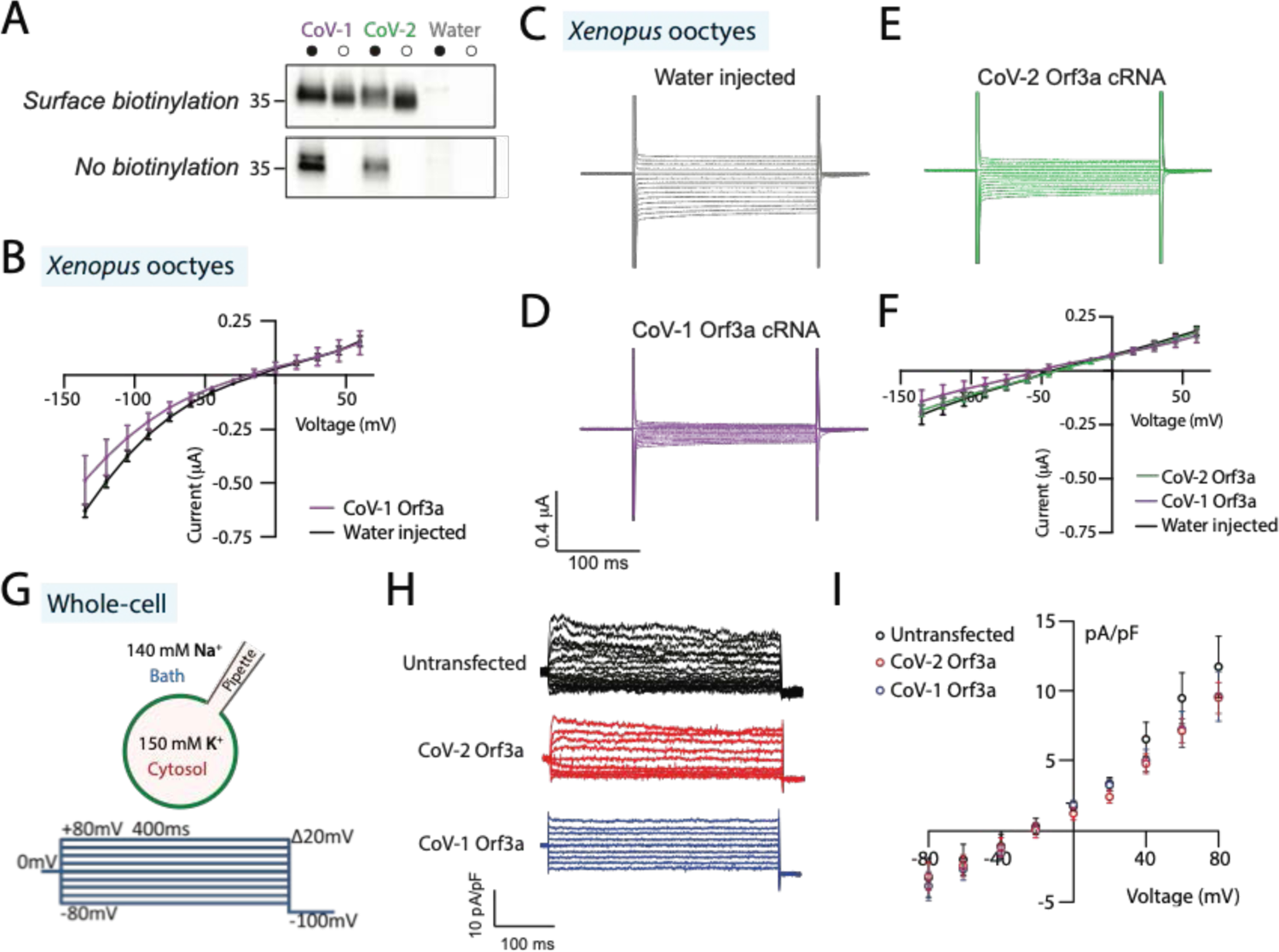
SARS-CoV-1 Orf3a is not a cationic ion channel at the plasma membrane of HEK293 cells and *Xenopus* oocytes. (**A**) Surface biotinylation experiments using *Xenopus* oocytes injected with SARS-CoV-1 (CoV-1) Orf3a_2x-STREP_, SARS-CoV-2 (CoV-2) Orf3a_2x-STREP_, or water demonstrates plasma membrane localization of Orf3a constructs used for two-electrode voltage clamp studies. Western blot detecting Orf3a_2x-STREP_, which migrates at ∼35 kDa. Total cell lysate was used as an input (filled circles). CoV-1 and CoV-2 Orf3a *Xenopus* oocytes surface biotinylation was detected when exposed to the biotin probe (open circles, surface biotinylation versus no biotinylation). (**B**) I-V relationship for water injected (black, n=7) or CoV Orf3a_2x-STREP_ mRNA injected (purple, n=7) *Xenopus* oocytes following protocol described in (Figure 2J-K). Water injected I-V trace is the same trace as in Figure 2L. (C-F) CoV-2 3a does not elicit a current in *Xenopus* oocytes in ND-96 external solution (Methods). Representative current traces from *Xenopus* oocytes injected with (**C**) water, (**D**) CoV-1 Orf3a_2x-STREP_ or (**E**) CoV-2 Orf3a_2x-STREP_ mRNA (20 μg). Recordings are done in ND-96 solution (96 mM NaCl) following a voltage protocol that recapitulates published methods. (F) I-V relationship for water injected (black, n=7), CoV-1 Orf3a (purple, n=7), or CoV-2 Orf3a (green, n=7) following protocol described in (C-E). (**G-I**) Neither CoV-1 nor CoV-2 Orf3a elicits a current at the surface of HEK293 cells. (**G**) Solutions and voltage step protocol used for whole-cell patch clamp experiments. (**H**) Representative current traces of HEK293 cells untransfected (black), or doxycycline-induced CoV-2 Orf3a_SNAP_ (red) or CoV-1 Orf3a_SNAP_ (blue) recorded using the voltage step protocol and solutions in (G). (**I**) I-V relationship for untransfected HEK293 cells (black, n=9), and cells doxycycline-induced to express CoV-2 Orf3a_SNAP_ (red, n=11) and CoV-1 Orf3a_SNAP_ (blue, n=9). Error is represented as SEM.

**Figure 3-figure supplement 1.**
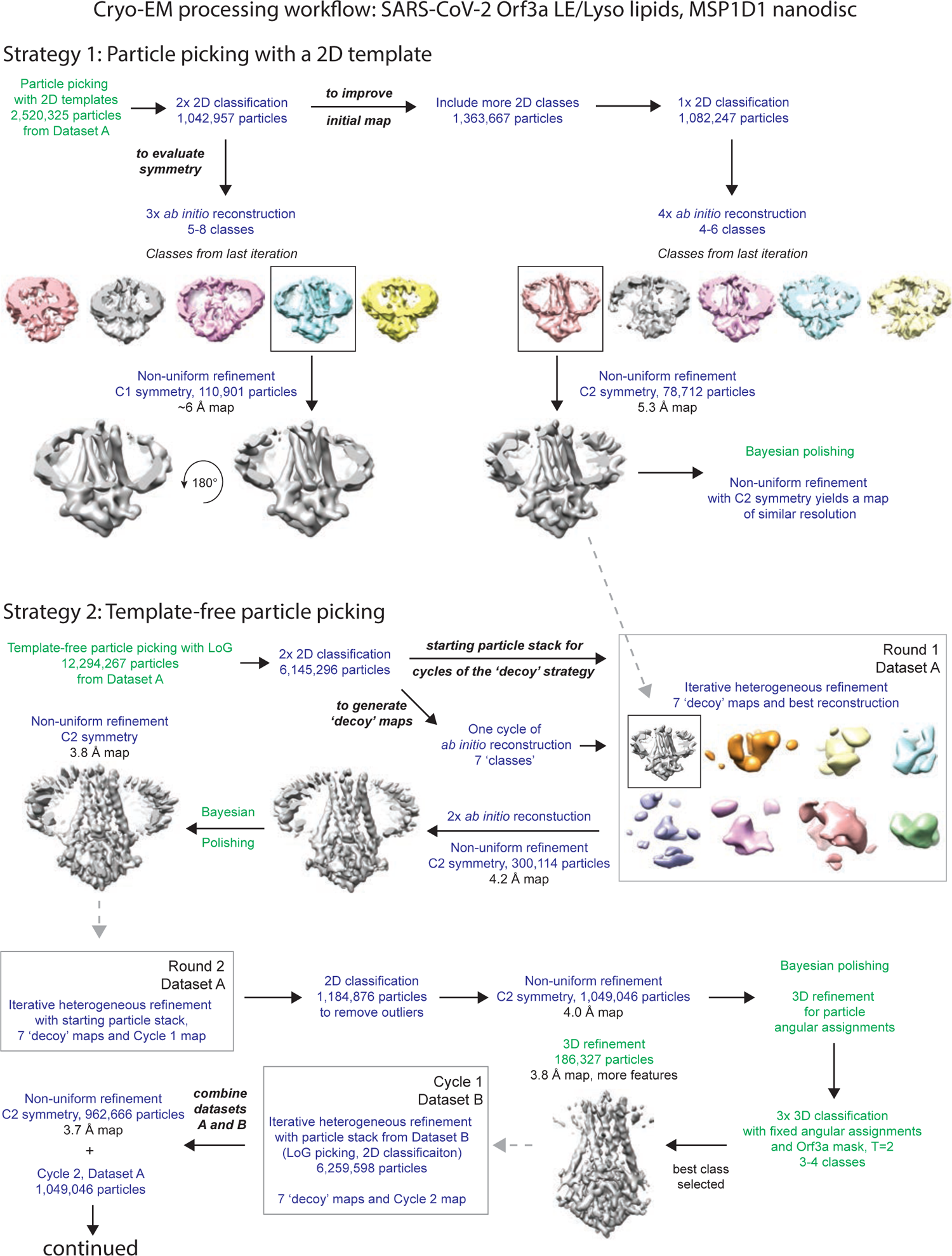
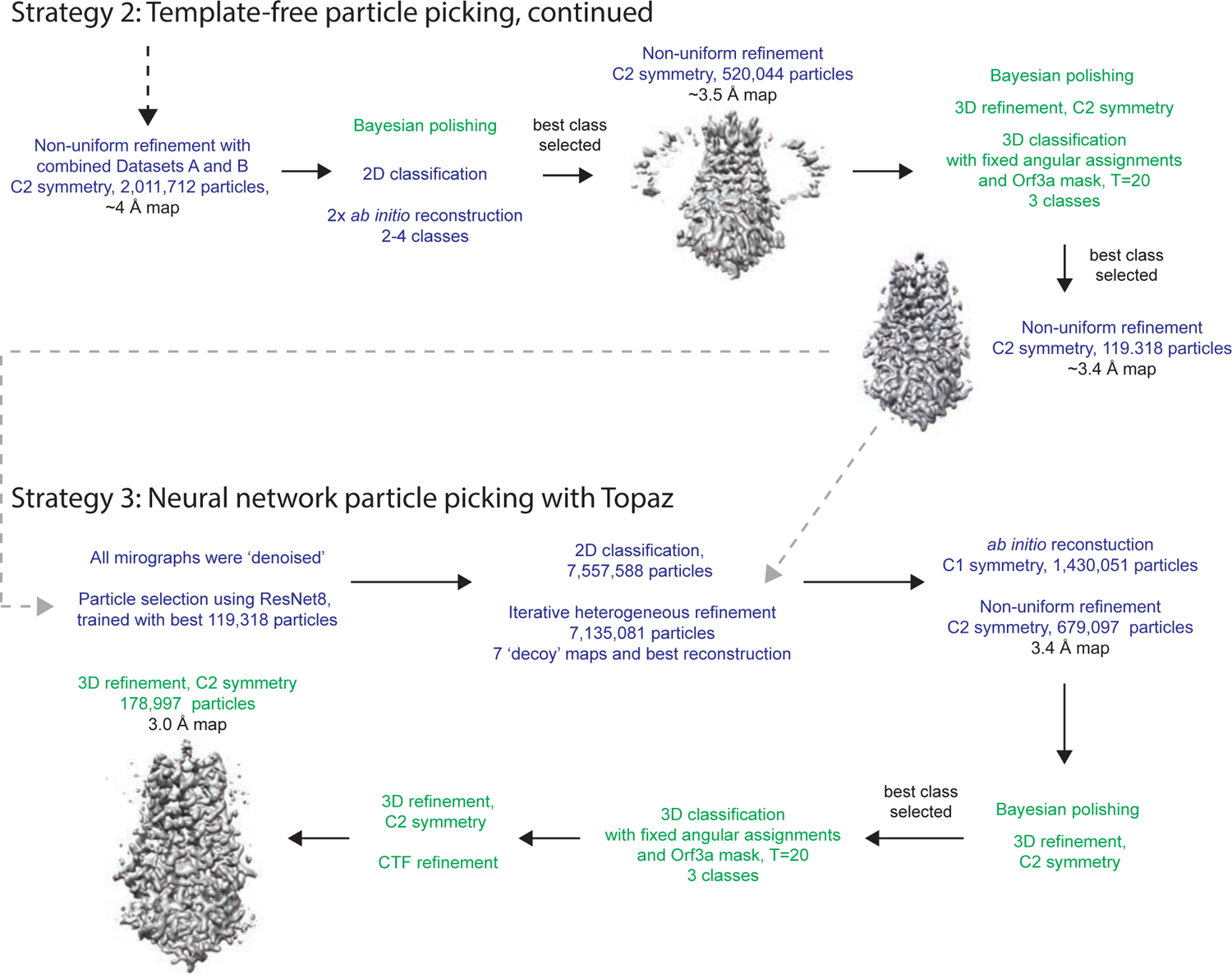
Cryo-EM data processing workflow for SARS-CoV-2 Orf3a reconstituted in LE/Lysosomal MSP1D1-containing nanodiscs. Text color denotes that the program Relion3.0 (green) or cryoSPARC3.0 (dark blue) was used for the step of the workflow.^61, 64^ Details are described in the Methods.

**Figure 3-figure supplement 2.**
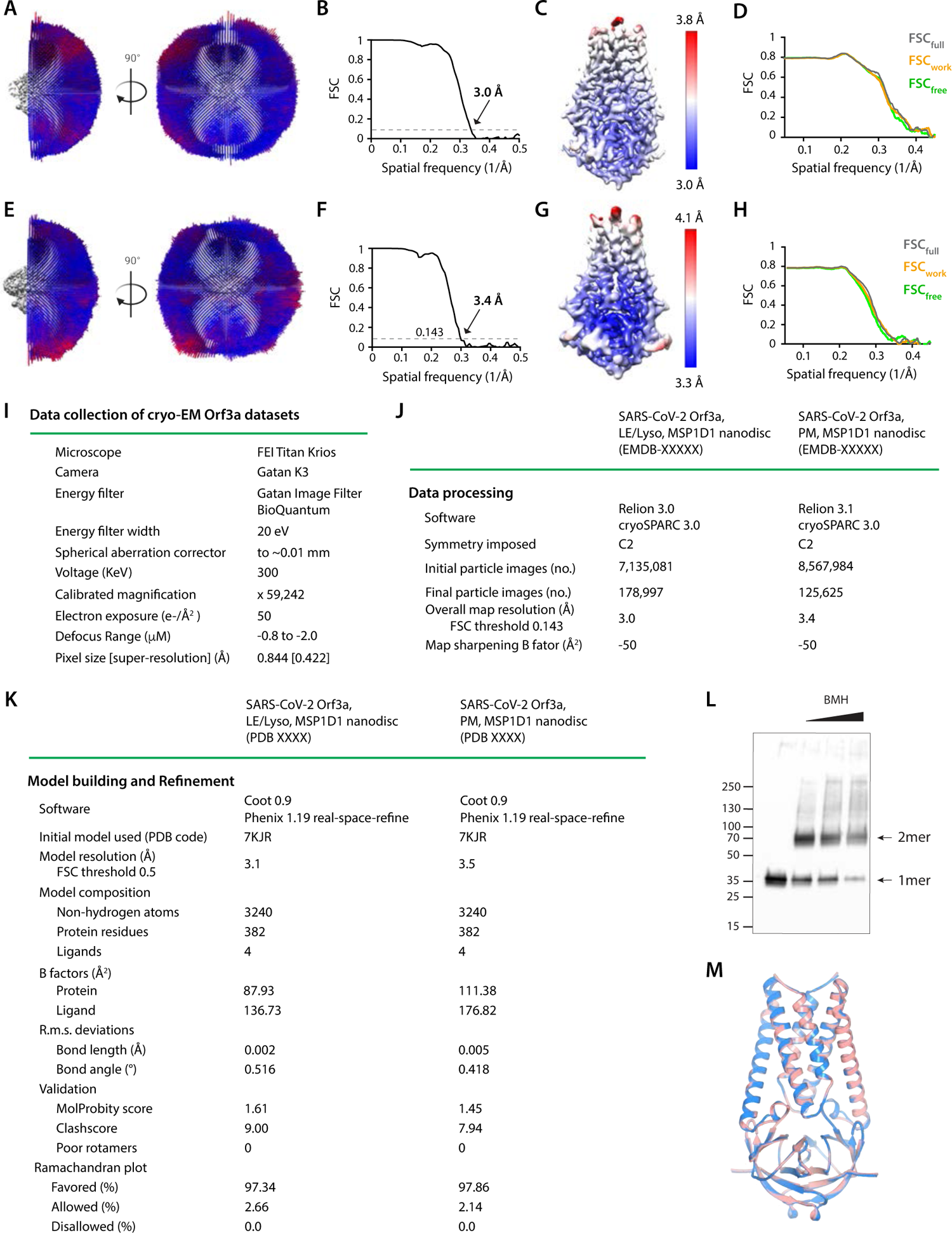
Structural determination of SARS-CoV-2 Orf3a LE/Lyso (**A-D**) or PM (**E-H**) MSP1D1 nanodiscs. (**A, E**) Angular orientation distribution of particles used in final reconstruction. The particle distribution is indicated by color shading, with blue to red representing low and high numbers of particles. (**B, F**) FSC curve of the final 3D reconstruction. The resolution is 3.0 Å (**B**) or 3.4 Å (**F**) at the FSC cutoff of 0.143 (dotted line). (**C, G**) Local resolution of the map was estimated using Relion and is colored as indicated. (D, H) Model validation. Comparison of the FSC curves between the model and half map 1 (FSC_work_), model and half map 2 (FSC_free_) and model and full map (FSC_full_). (**I-K**) Table of data collection and model statistics for the SARS-CoV-2 Orf3a LE/Lyso or PM MSP1D1 nanodisc structures. (**L**) Crosslinking of SARS-CoV-2 Orf3a from isolated HEK293 cellular membrane to assess its oligomeric state. A band of the approximate molecular weight of a dimer (2 mer) appears with the addition of bismaleimidohexane (BMH). (**M**) Superposition of the SARS-CoV-2 Orf3a LE/Lyso MSP1D1 (pink) and PM (blue) structures.

**Figure 3-figure supplement 3.**
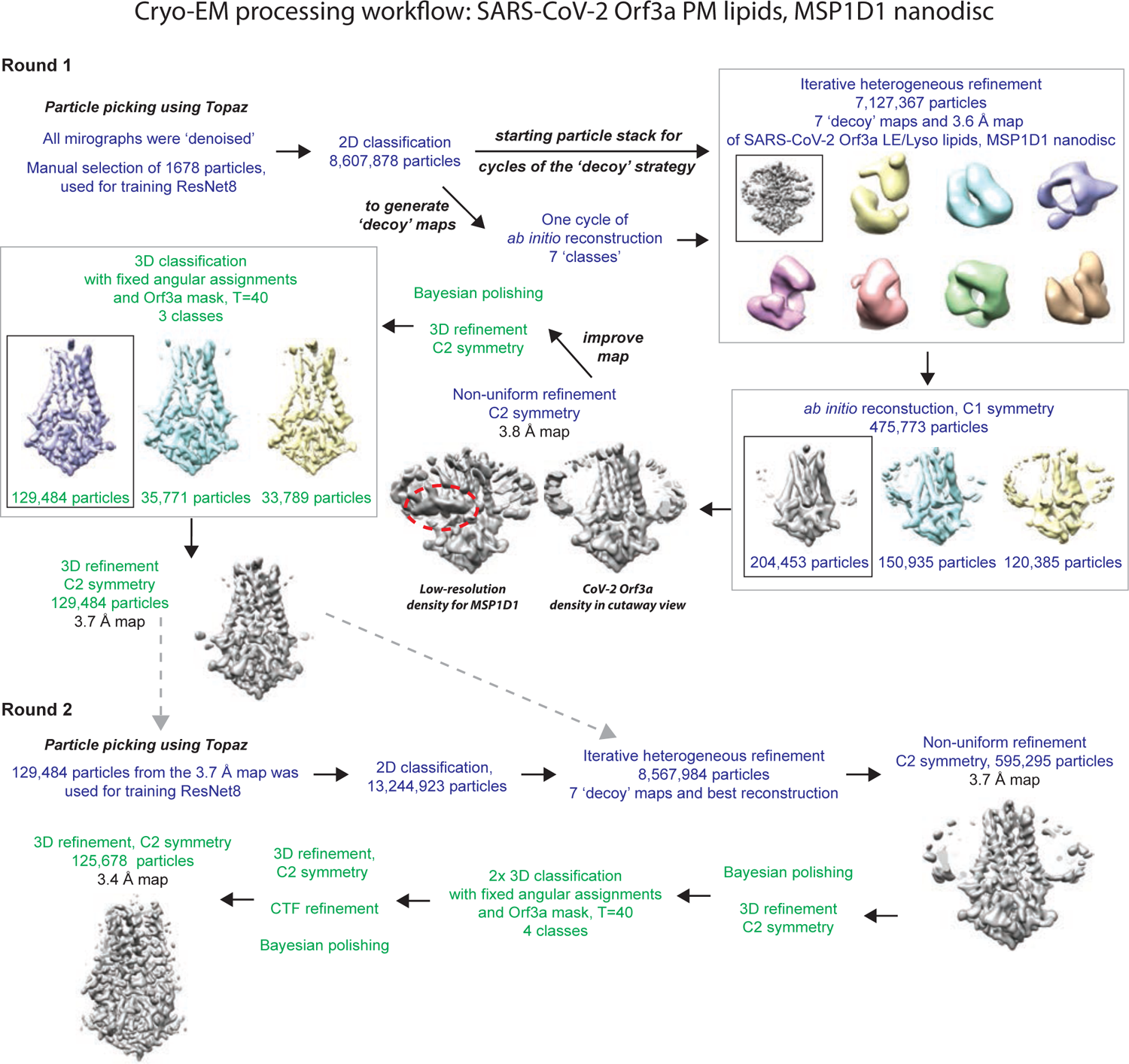
Cryo-EM data processing workflow for SARS-CoV-2 Orf3a reconstituted in PM MSP1D1-containing nanodiscs. Text color denotes that the program Relion3.1 (green) or cryoSPARC3.0 (dark blue) used.^61, 64^ See Methods. Low-resolution density for MSP1D1 is visible in maps of CoV-2 Orf3a but does not resolve to high-resolution (red circle).

**Figure 3-figure supplement 4.**
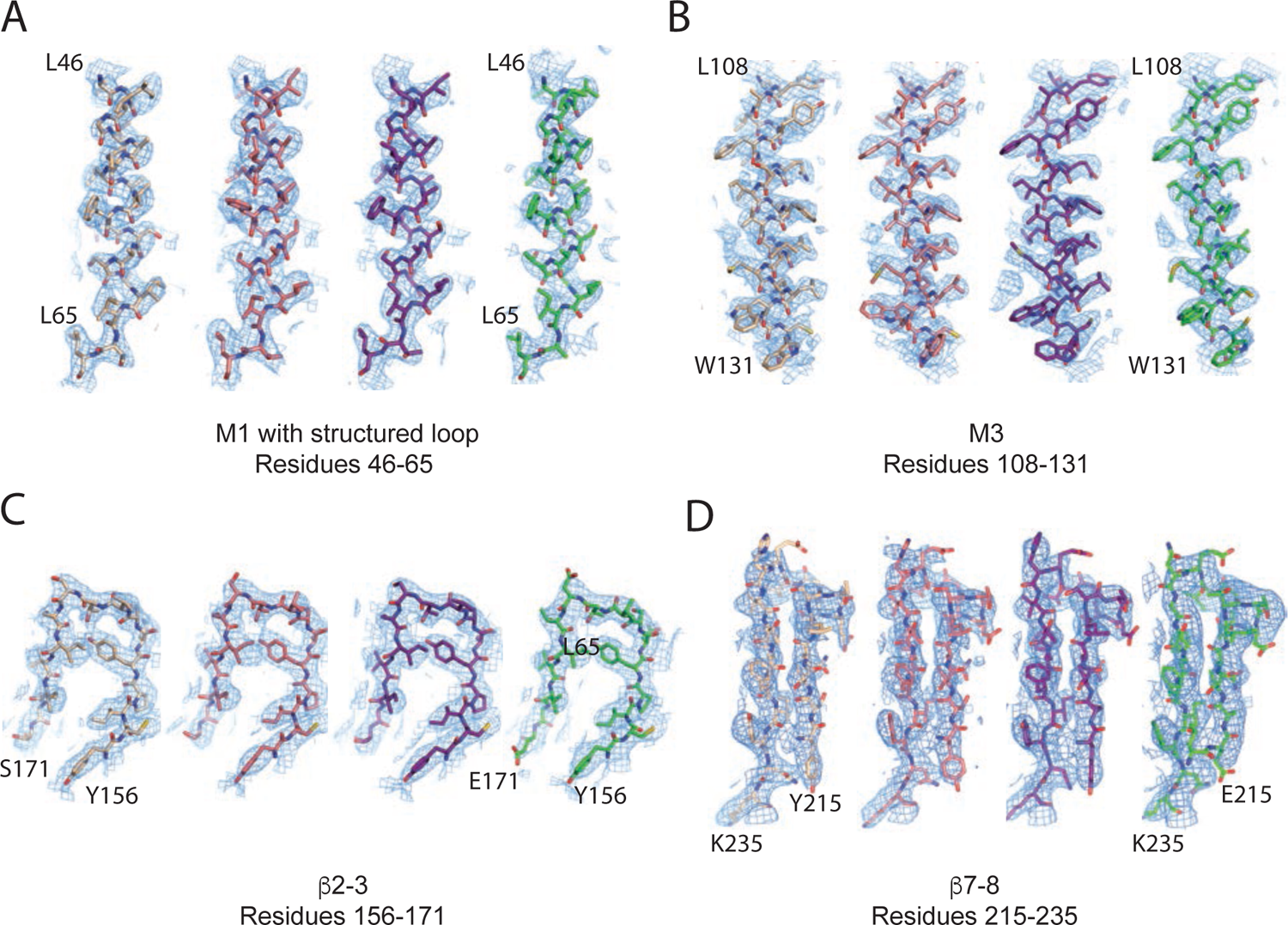
Representative cryo-EM density for SARS-CoV-2 Orf3a and SARS-CoV-1 Orf3a structures. (**A-D**) Four representative areas of cryo-EM density (blue mesh) from the four Orf3a datasets with structures represented as sticks and colored as follows: SARS-CoV-2 Orf3a LE/Lyso MSP1D1-containing nanodisc (tan), SARS-CoV-2 Orf3a PM MSP1D1-containing nanodisc (pink), SARS-CoV-2 LE/Lyso Saposin A-containing nanodisc (dark purple), SARS-CoV-1 Orf3a LE/Lyso MSP1D1-containing nanodisc (green).

**Figure 4-figure supplement 1.**
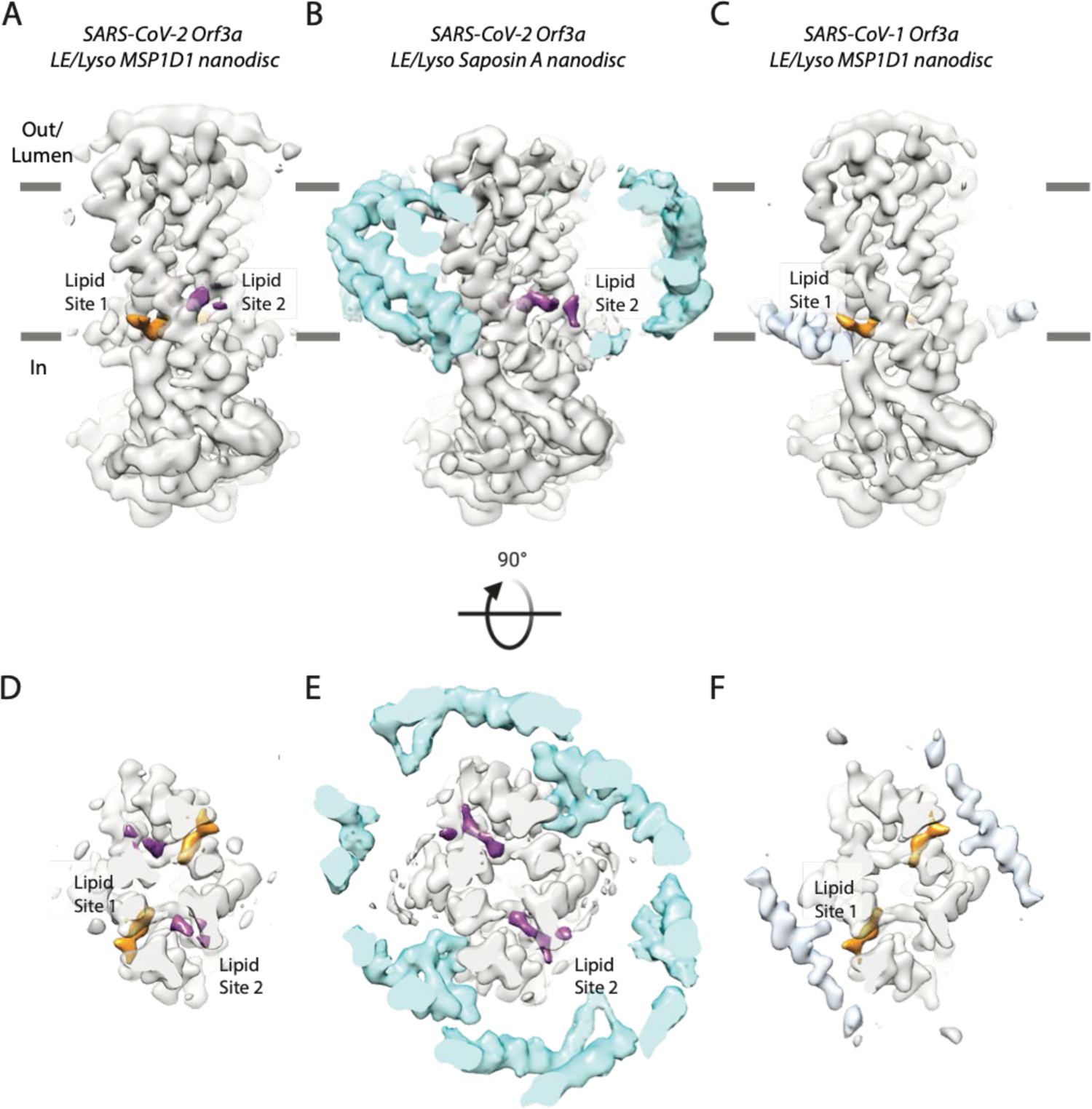
Comparison of lipid densities between (**A, D**) SARS-CoV2 Orf3a LE/Lyso MSP1D1-containing nanodiscs (**B, E**) SARS-CoV2 Orf3a LE/Lyso Saposin A-containing nanodiscs and (**C, F**) SARS-CoV-1 Orf3a LE/Lyso MSP1D1-containing nanodiscs. Side views (**A-C**) and cutaway views (**D-F**) from the extracellular/luminal side of the membrane. Density for Lipid Site 1 (orange) and Lipid Site 2 (purple) highlights distinct binding between structures. Saposin A (**B, E**) and MSP1D1 (**C, F**) densities are depicted in cyan and light blue, respectively.

**Figure 4-figure supplement 2.**
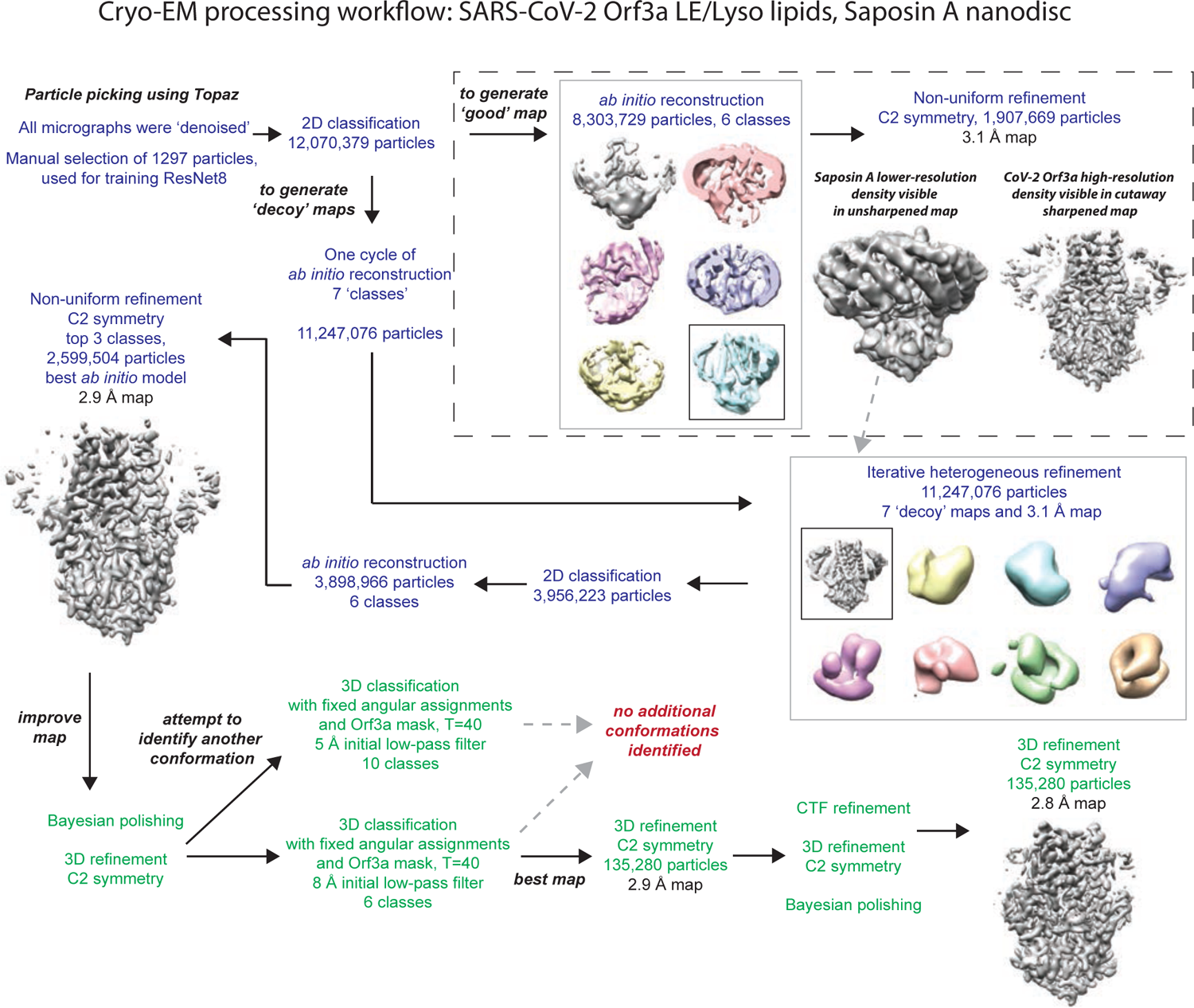
Cryo-EM data processing workflow for SARS-CoV-2 Orf3a reconstituted in LE/Lyso Saposin A-containing nanodiscs. Text color denotes that the program Relion3.1 (green) or cryoSPARC3.0 (dark blue).^61, 64^ See Methods. The initial map generated for iterative heterogeneous refinement (dotted box) displayed both higher-resolution density for CoV-2 Orf3a (gray, sharpened map) and lower-resolution Saposin A molecules surrounding SARS-CoV-2 Orf3a (gray, unsharpened map).

**Figure 4-figure supplement 3.**
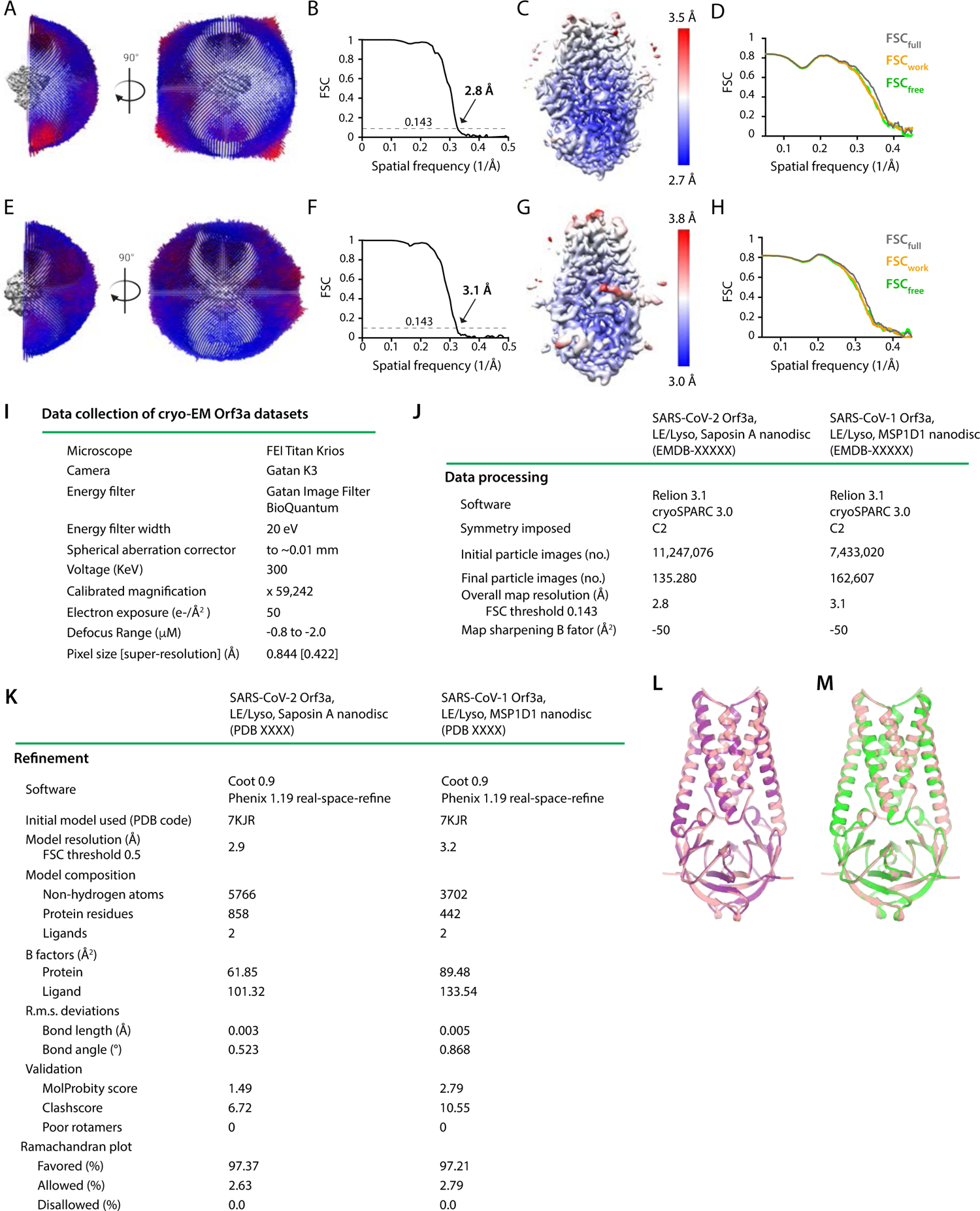
Structural determination of SARS-CoV-2 Orf3a LE/Lyso Saposin A nanodisc (**A-D**) or SARS-CoV-1 LE/Lyso MSP1D1 nanodisc (**E-H**). (**A, E**) Angular orientation distribution of particles used in final reconstruction. The particle distribution is indicated by color shading, with blue to red representing low and high numbers of particles. (**B, F**) FSC curve of the final 3D reconstruction. The resolution is 2.8 Å (B) or 3.1 Å (**F**) at the FSC cutoff of 0.143 (dotted line). (C, G) Local resolution of the map was estimated using Relion and is colored as indicated. (**D, H**) Model validation. Comparison of the FSC curves between the model and half map 1 (FSC_work_), model and half map 2 (FSC_free_) and model and full map (FSC_full_). (**I-K**) Table of data collection and model statistics for the SARS-CoV-2 Orf3a LE/Lyso Saposin A or SARS-CoV-1 Orf3a LE/Lyso MSP1D1 nanodisc structures. (**L**) Superposition of the SARS-CoV-2 Orf3a LE/Lyso MSP1D1 (pink) and SARS-CoV-2 Orf3a LE/Lyso Saposin A (purple) structures. (**M**) Superposition of the SARS-CoV-2 Orf3a (pink) and SARS-CoV-1 Orf3a (green) LE/Lyso MSP1D1 structures.

**Figure 4-figure supplement 4.**
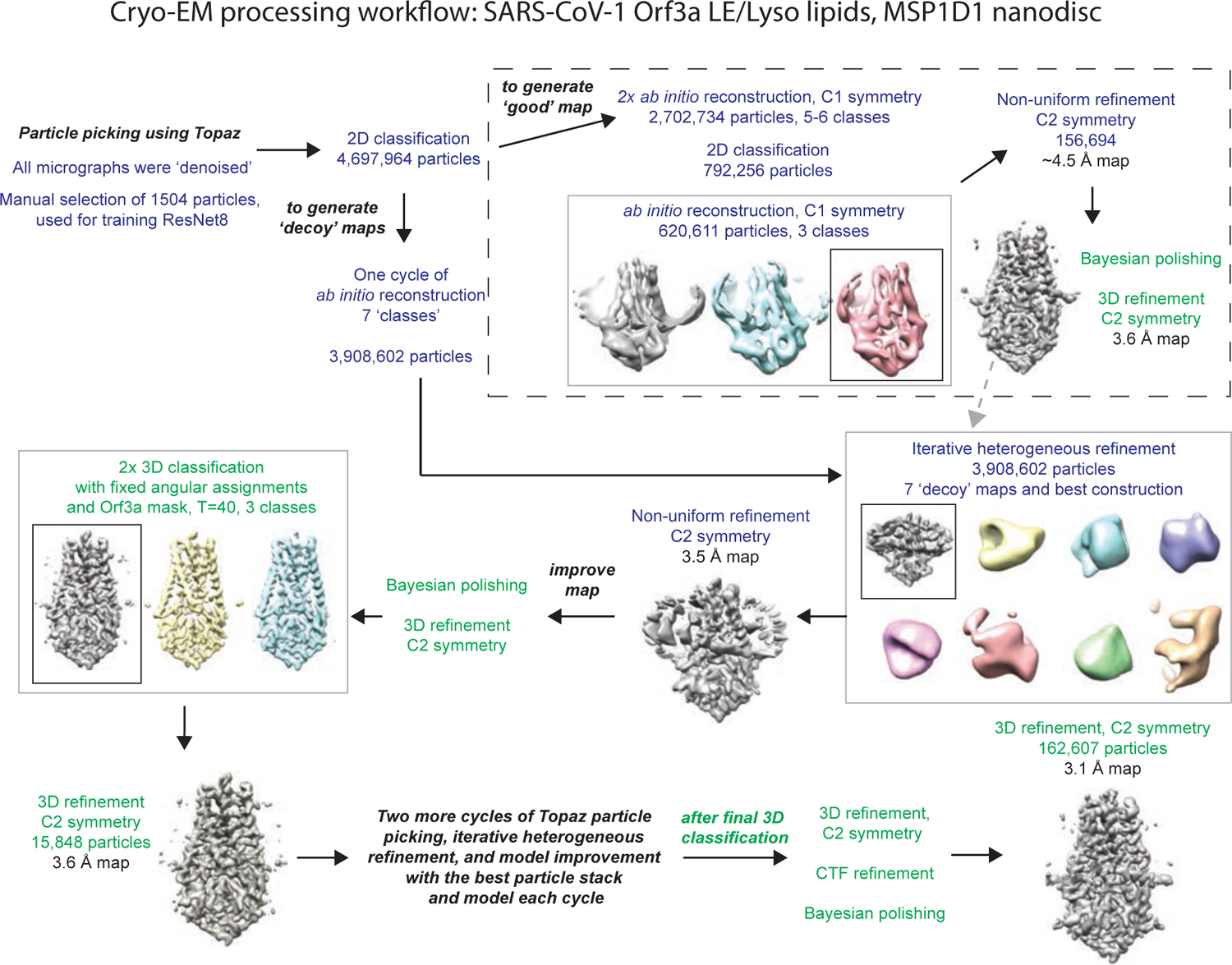
Cryo-EM data processing workflow for SARS-CoV-1 Orf3a reconstituted in LE/Lysosomal MSP1D1-containing nanodiscs. Text color denotes the program Relion3.1 (green) or cryoSPARC3.0 (dark blue). ^61, 64^ See Methods.

**Figure 4-figure supplement 5.**
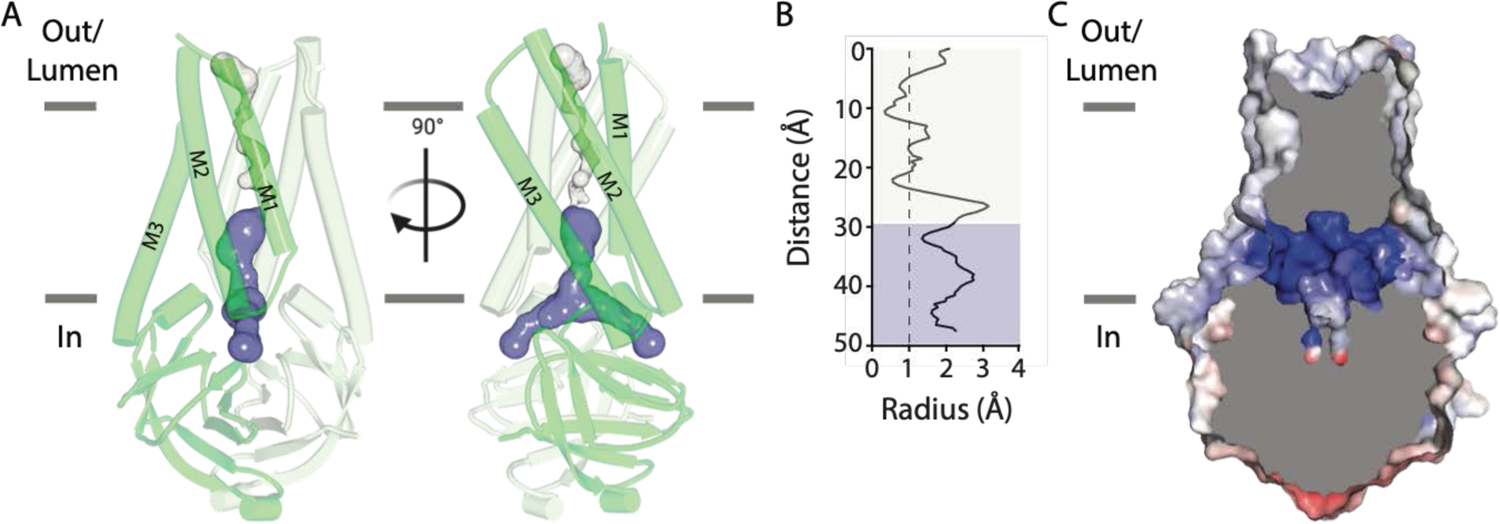
A similar narrow cavity is detected in the TM region of SARS-CoV-1 Orf3a. (**A**) Two representative side views of SARS-CoV-1 (CoV-1) Orf3a in LE/Lyso MSP1D1-containing nanodiscs highlighting two subunits (dark and light pink). Inspection of the transmembrane region for a pore, depicted as the minimal radial distance from its center to the nearest van der Waals protein contact (HOLE program).^73^ A region too narrow to conduct ions (white) and an aqueous vestibule (dark blue) are highlighted. (**B**) Radius of the ion pore (from A) as a function of the distance along the ion pathway. Dashed lines indicate the minimal radius that would permit a dehydrated cation. Blue and white colors follow HOLE diagram of A. (**C**) The CoV-1 Orf3a molecular surface is colored according to the electrostatic potential (APBS program).^74^ Coloring: blue, positive (+10 kT/e) and red, negative (−10 kT/e). Cutaway view looking into the aqueous vestibule.

**Figure 5-figure supplement 1.**
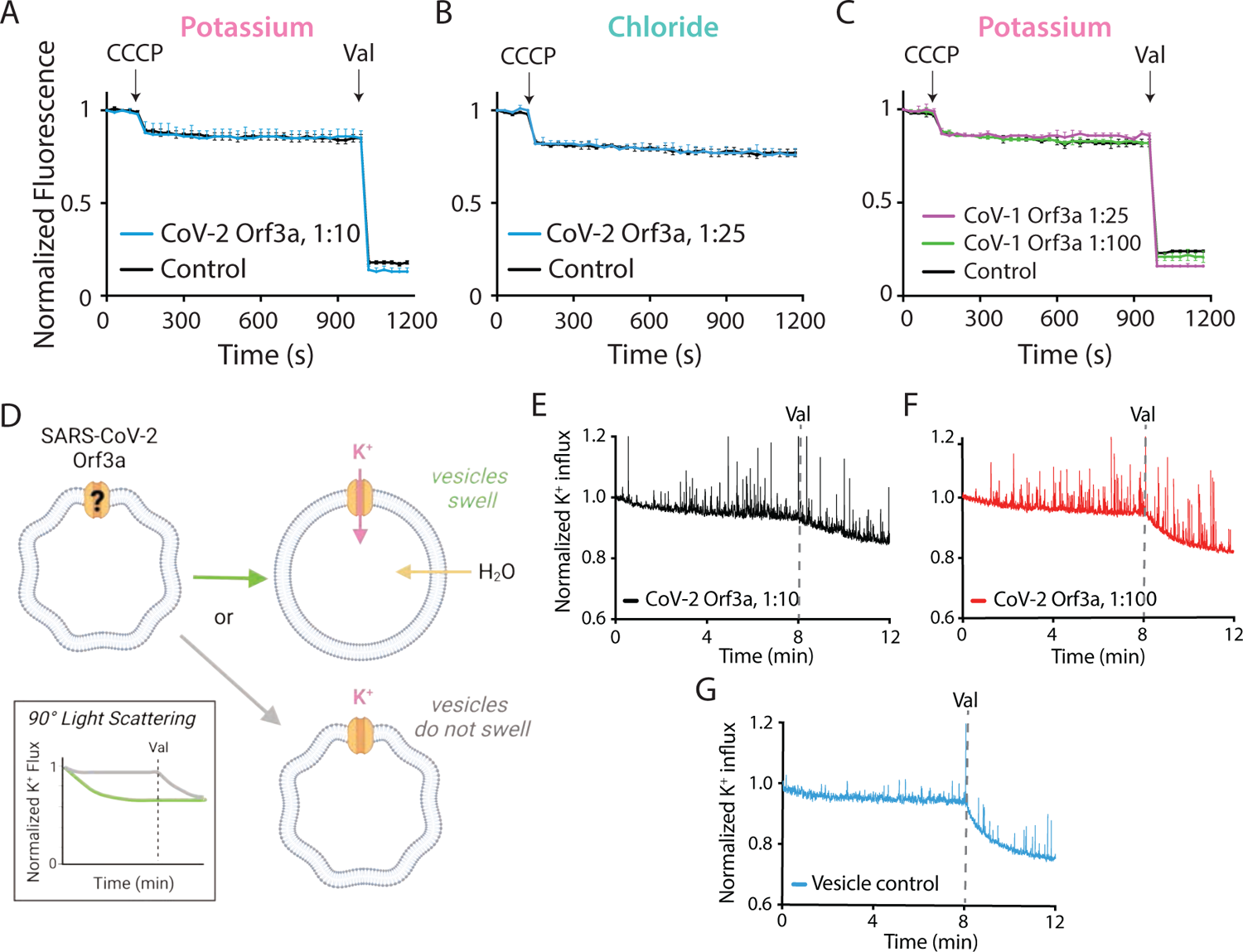
Characterization of vesicle-reconstituted SARS-CoV-1 and SARS-CoV-2 Orf3a at low and high protein ratios. (**A-B**) K^+^ (A) or Cl^-^ (**B**) flux is not observed in SARS-CoV-2 Orf3a-reconstituted vesicles (blue) as compared with the empty vesicle control (black, n=3) using 1:10 (n=3) (A) or 1:25 (n=3) (B) ratios of protein to lipids. Error is represented as SEM. (**C**) K^+^ flux is not observed in SARS-CoV-1 Orf3a-reconstituted vesicles (blue) as compared with the empty vesicle control (black, n=3) using a 1:25 (purple, n=3) or 1:100 (green, n=3) ratio of protein to lipids. Error is represented as SEM. (**D**) Schematic of 90° light-scattering K^+^ flux assay.^36, 37^ SARS-CoV-2 Orf3a vesicles reconstituted in 200 mM K-glutamate are diluted into a hypertonic buffer containing 260 mM K-thiocyanate, resulting in vesicle shrinkage. If SARS-CoV-2 Orf3a is a K^+^-selective viroporin, then the asymmetrical K^+^ concentration should drive K^+^ influx, leading to water absorption, vesicle swelling (green arrow) and a reduction of 90° light-scattering (green line, inset). If SARS-CoV-2 Orf3a is not a K^+^-selective viroporin, then vesicles will not swell (gray arrow) and no change in 90° light-scattering should be observed (gray line, inset). The addition of valinomycin (Val) leads to vesicle swelling and reduction of 90° light-scattering should be observed for all vesicles in the sample. Created with Biorender.com. (**E-G**) No difference in normalized K^+^ influx is observed among SARS-CoV-2 Orf3a reconstituted using 1:10 (E, black) or 1:100 (F, red) ratio of protein to lipids, and control vesicles (G, blue).

**Figure 5-figure supplement 2.**
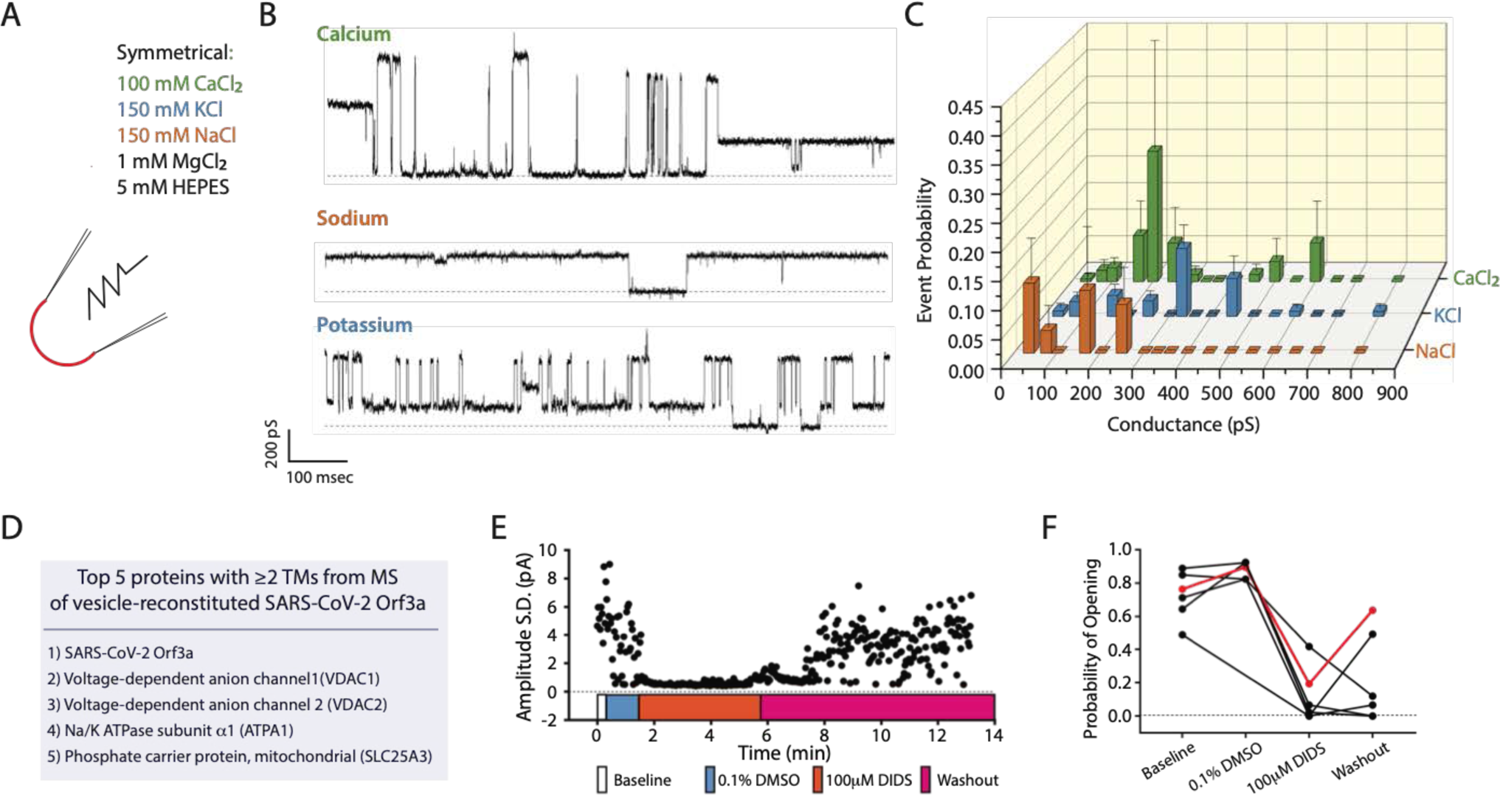
Currents recorded from SARS-CoV-2 Orf3a containing-vesicles reconstituted at a high protein to lipid ratio likely result from transient membrane leakiness and/or contamination by *bona fide* ion channels. (**A**) Symmetrical recording solutions with CaCl_2_ (green), KCl (blue) or NaCl (orange) used for proteoliposome patch clamp experiments. (**B**) Multiple K^+^, Na^+^ and Ca^2+^ conductances species are observed by proteoliposome patch clamp with SARS-CoV-2 Orf3a-containing vesicles reconstituted at a 1:10, but not 1:100 (Figure 5E), ratio of protein to lipid. (**C**) Probability of observing an open event in a SARS-CoV-2 Orf3a reconstituted proteoliposome patch with vesicles reconstituted at a 1:10 ratio of protein to lipid. NaCl n = 18, KCl n = 105, and CaCl_2_ n = 22. Error is presented as SEM. (B). (**D**) Table of the top 5 proteins with 2TM helices or greater from mass spectrometry analysis of SARS-CoV-2 Orf3a containing vesicles reconstituted at a 1:10 protein to lipid ratio. (**E**) Addition of 100 μM 4,4’-Diisothiocyano-2,2’-stilbenedisulfonic Acid (DIDS; orange trace), a blocker of VDAC channels, eliminates the Ca^2+^ currents observed with proteoliposome patches-reconstituted CoV-2 Orf3a, whereas DMSO has no effect (blue). (**F**) Open probability of proteoliposome patches-reconstituted CoV-2 Orf3a sample following protocol from (E).

**Figure 6-figure supplement 1.**
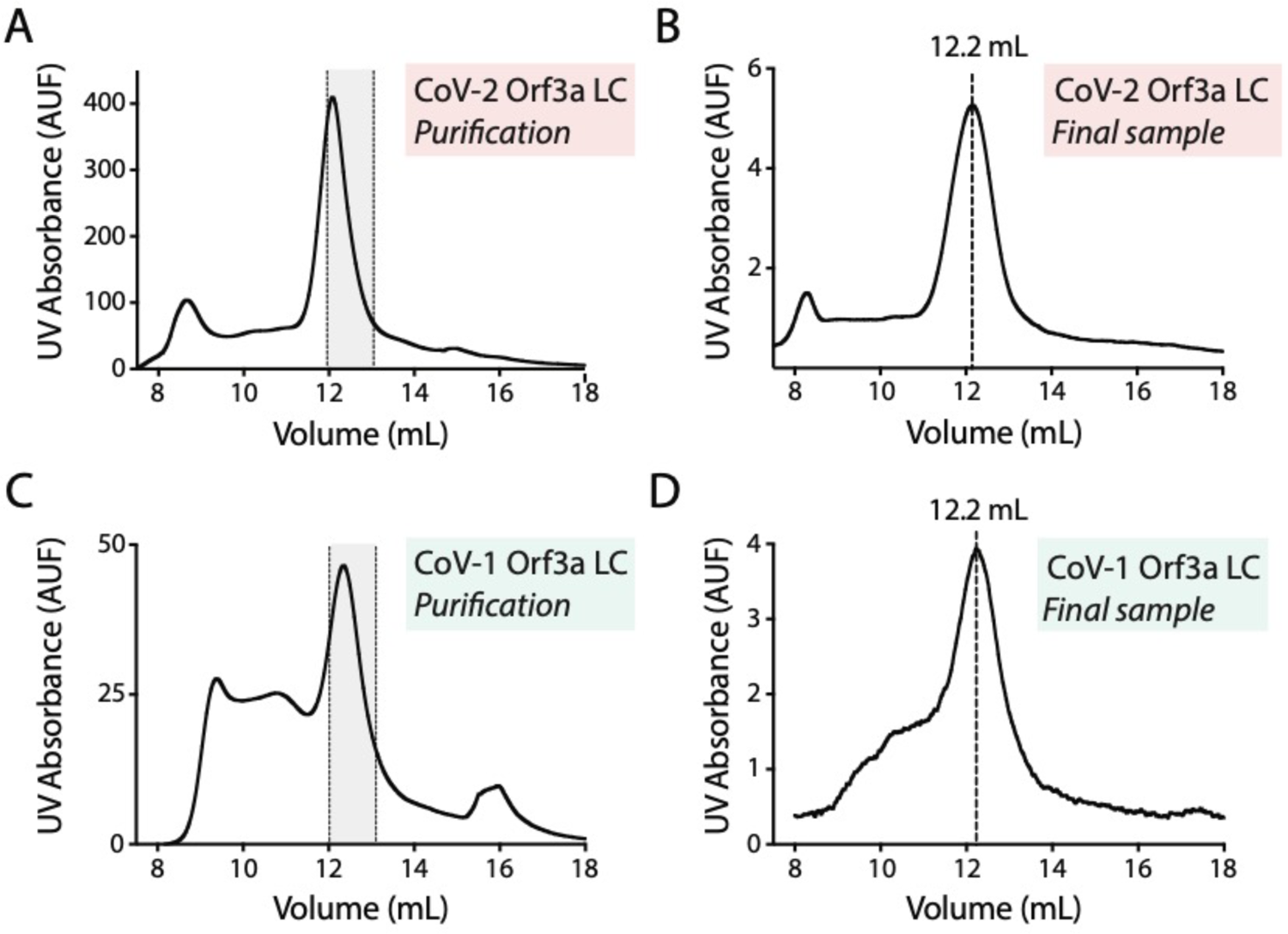
Purification of SARS-CoV-1 and SARS CoV-2 Orf3a loop chimeras. (**A, C**) Gel filtration traces from (**A**) SARS-CoV-2 Orf3a loop chimera (CoV-2 Orf3a LC) and (**C**) SARS-CoV-1 Orf3a loop chimera (CoV-1 Orf3a LC) after elution from StrepTactinXT column. Collected peak fraction is highlighted in gray. (**B, D**) Gel filtration traces of final samples of (**B**) CoV-2 Orf3a LC and (**D**) CoV-1 Orf3a LC used for the co-IP experiments (Figure 6G-H).

## Notes

### Competing Interest Statement

The authors have declared no competing interest.

